# Conserved noncoding transcription and core promoter regulatory code in early *Drosophila* development

**DOI:** 10.1101/156596

**Authors:** Philippe J. Batut, Thomas R. Gingeras

## Abstract

Multicellular development is largely determined by transcriptional regulatory programs that orchestrate the expression of thousands of protein-coding and noncoding genes. To decipher the genomic regulatory code that specifies these programs, and to investigate globally the developmental relevance of noncoding transcription, we profiled genome-wide promoter activity throughout embryonic development in 5 *Drosophila* species. We show that core promoters, generally not thought to play a significant regulatory role, in fact impart broad restrictions on the developmental timing of gene expression on a genome-wide scale. We propose a hierarchical model of transcriptional regulation during development in which core promoters define broad windows of opportunity for expression, by defining a limited range of transcription factors from which they are able to receive regulatory inputs. This two-tiered mechanism globally orchestrates developmental gene expression, including noncoding transcription on a scale that defies our current understanding of ontogenesis. Indeed, noncoding transcripts are far more prevalent than ever reported before, with ∼4,000 long noncoding RNAs expressed during embryogenesis. Over 1,500 are functionally conserved throughout the *melanogaster* subgroup, and hundreds are under strong purifying selection. Overall, this work introduces a hierarchical model for the developmental regulation of transcription, and reveals the central role of noncoding transcription in animal development.

## Introduction

Development in metazoans is orchestrated by complex gene regulatory programs encoded in the sequence of the genome^1-4^. The expression of thousands of protein-coding and noncoding genes, in precise spatiotemporal patterns, progressively refines the organization of embryonic structures and specifies the differentiation of specialized cell types. In *Drosophila,* many of the master genes governing early development^1,2^ encode regulators of transcription^5-8^, and transcriptional regulation largely accounts for embryo patterning^5,9^.

The regulatory code that specifies these programs, however, remains poorly understood. Sequences that bind transcriptional activators and repressors, known as enhancers^10-16^, are generally thought to be the primary determinants of gene expression specificity. Sequences that serve as docking sites for the basal transcription machinery, the core promoters^17^, are usually assumed to be structural elements that contribute little or no regulatory information.

Indeed, core promoters contain sequence motifs (*e.g.,* TATA boxes) that serve as a platform for RNA Pol II initiation at transcription start sites (TSSs), but are not by themselves sufficient to induce transcription^14,17^. Sequence-specific activators and repressors, collectively designated as transcription factors (TFs), bind to enhancers and foster the assembly of the basal transcription machinery at associated core promoters^14,17^. Beyond these general principles, the syntax of the code, and in particular the functional relationship between these two classes of elements, remains obscure. Interacting core promoter-enhancer pairs may be directly adjacent^12,16,18^, or may be located hundreds of kilobases apart in metazoan genomes^14,19^, and the rules enforcing specific interactions are unknown^14,17^. There is evidence that core promoters can influence the expression specificity of some genes^18,20^, but so far this has not been systematically studied in the context of development. Understanding the basis of transcriptional control requires parsing out these complex interactions.

In addition to delineating the rules of gene regulation, it is necessary to expand the concept of gene^21,22^ to include all the noncoding loci that may control developmental processes. Indeed, it has become increasingly clear that noncoding transcription is very prevalent in Eukaryotes^23-28^, and both genetic and biochemical studies have unambiguously established long noncoding RNAs (lncRNAs) as functional components of the cellular machinery^29-32^. Exhaustively annotating noncoding transcripts, and identifying those with biologically relevant functions, is crucial to our understanding of development.

Deciphering the meaning of regulatory sequences, or assessing the biological relevance of lncRNA genes, are daunting tasks that require innovative strategies. The use of genome-wide functional assays in a phylogenetic framework is a powerful and general approach to such questions^33-36^. Indeed, direct measurements of complex genome functions in multiple species provide a sort of genomic Rosetta Stone from which the underlying code can begin to be parsed out.

Here, we used high-throughput TSS mapping in tightly resolved time series to establish genome-wide promoter activity profiles throughout embryonic development in 5 *Drosophila* species spanning 2550 million years (MY) of evolution. Combining TSS identification at single-nucleotide resolution^37^ with quantitative measurements of developmental expression patterns, we uncovered unique features of expression timing and core promoter structure to generate novel insights into transcriptional regulation.

We report that distinct types of core promoters are selectively active in three broad phases of embryonic development: specific combinations of core motifs mediate transcription during Early, Intermediate and Late embryogenesis. Each individual class of core promoters is functionally associated with distinct sets of transcription factors. We propose a two-tier model of transcriptional control in which core promoters and enhancers mediate, respectively, coarse-grained and fine-grained developmental regulation.

We also show that noncoding transcription is far more widespread than anticipated in *Drosophila,* with 3,973 promoters driving the expression of lncRNAs during embryogenesis. The analysis of these core promoters, most of which are currently unannotated, shows that they largely share the structural and functional properties of their counterparts at protein-coding genes. Through the analysis of their fine structure and sequence conservation, we demonstrate that evolutionarily conserved lncRNAs promoters are under strong purifying selection at the levels of primary sequence and expression specificity. We functionally characterize the *schnurri-like RNA* (slr) locus, which expresses a lncRNA in a highly conserved spatiotemporal pattern suggestive of a role in early dorsoventral patterning.

In summary, these results uncover a major active role for core promoters in regulating transcription, by defining windows of opportunity for activation by enhancer sequences. They also reveal a vastly underappreciated aspect of developmental transcriptomes, by showing that noncoding transcription is extremely prevalent, tightly regulated and, crucially, deeply conserved.

## Results

### Global multispecies profiling of developmental promoter activity

To explore the genome-wide dynamics of transcriptional regulation and their evolution, we generated developmental transcriptome profiles at both very high temporal resolution (1 hour) and high sequence coverage (137-180 million read pairs per species) for 5 *Drosophila* species spanning 25-50 million years of evolution: *D. melanogaster, D. simulans, D. erecta, D. ananassae* and *D. pseudoobscura* (Fig. 1A; total of 120 samples). We focused on embryonic development, a crucial period during which the body plan is established and all larval organs are generated. This data allows direct, genome-wide comparisons of promoter activity in a phylogenetic framework (Fig. 1B).

**Figure 1:**
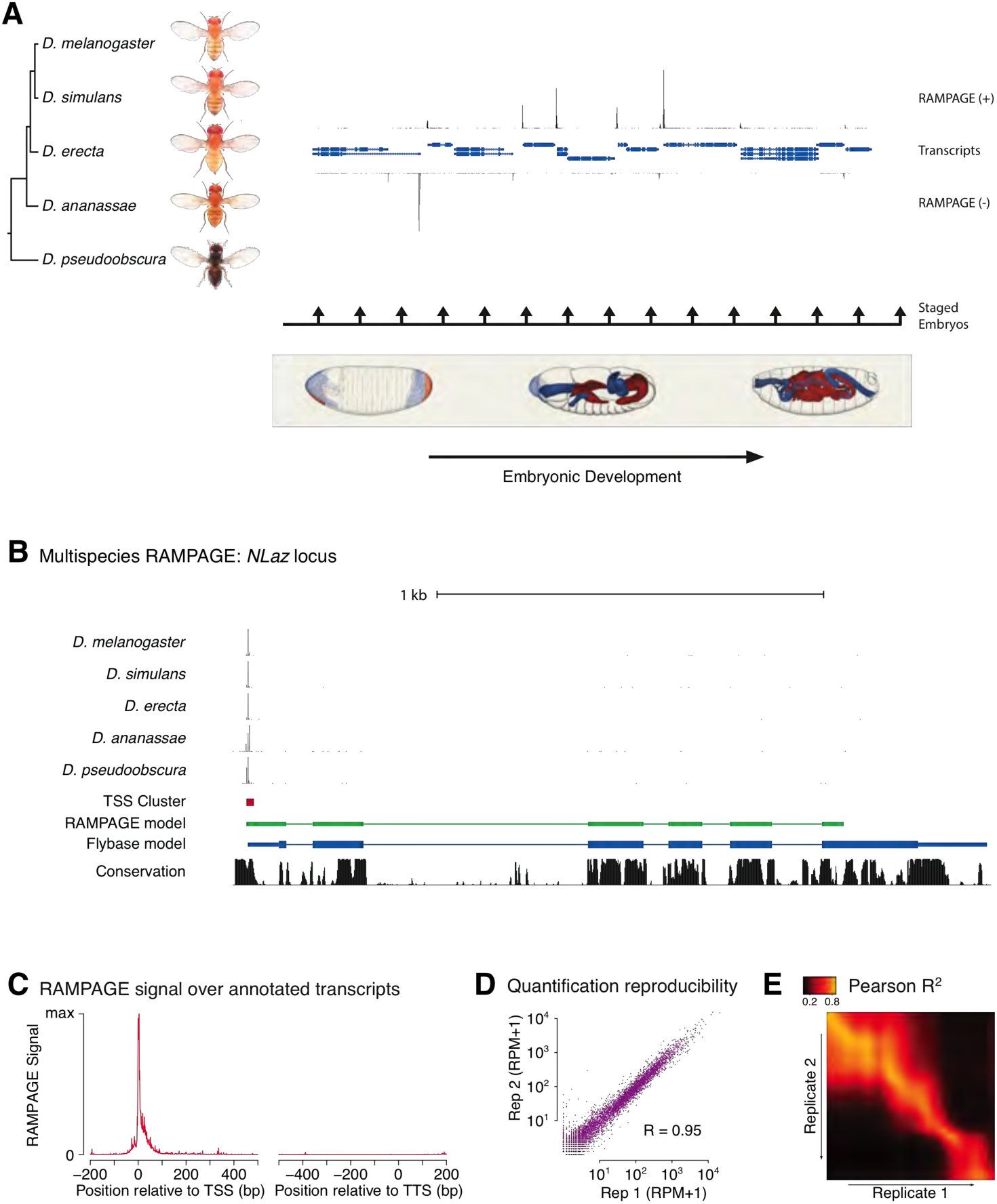
Comparative profiling of embryonic promoter activity. (a) Genome-wide TSS usage maps were generated by RAMPAGE assays for developmental series in 5 species (central panel, locus chr2L:1151500-1185900). Blue symbols: FlyBase transcript annotations; black density plot: density of RAMPAGE read 5’ ends on the positive strand; grey density plot: read 5’ ends on the negative strand (inverted y-axis). Fly photographs (N. Gompel) and embryo drawings (V. Hartenstein, CSHL Press 1993) reproduced with permission from FlyBase. (b) Distribution of RAMPAGE signal over the *NLaz* gene in 5 *Drosophila* species. From top: RAMPAGE signal intensity tracks, *D. melanogaster* TSC, transcript model inferred from RAMPAGE data, transcript model from FlyBase, sequence conservation (phastCons). For visualization of *non-melanogaster* data, sequencing reads were mapped to the appropriate genomes and projected onto orthologous *D. melanogaster* positions based on whole-genome alignments. (c) Metaprofile of RAMPAGE read 5’ ends over FlyBase-annotated mature transcripts (introns excluded). (d) Reproducibility of RAMPAGE signal quantification for individual TSCs (n=9,299) for two biological replicates of the first *D. melanogaster* time point (0-1h). TSCs with no signal in either replicate were excluded. (e) Correlation matrix for the *D. melanogaster* time series biological replicates. We plotted the correlation of TSC expression (n=24,832 TSCs) after smoothing and alignment of the time series.

In order to map transcription start sites (TSSs) with single-base resolution and accurately measure the activity of individual promoters, we used a high-fidelity method called RAMPAGE^37^ based on high-throughput sequencing of 5’-complete complementary DNA (cDNA) molecules. It offers unprecedented specificity and detection sensitivity for TSSs, and its multiplexing capabilities allow for the seamless acquisition of high-resolution developmental time series^37^. Since eukaryotic promoters often allow productive transcription initiation from multiple positions^37-40^, we use a dedicated peak-calling algorithm to group neighboring TSSs into TSS clusters (TSCs) corresponding to individual promoters^37,41^.

For each species, we identified 2.2x10^4^ to 2.7x10^4^ high-confidence TSCs. The narrow distribution of raw RAMPAGE signal (Fig. 1C & S1) and of TSCs (Fig. S2) over annotated loci confirms our very high specificity for true TSSs. The quantification of promoter expression levels is highly reproducible across biological replicates (Fig. 1D-E). Importantly, paired-end sequencing of cDNAs allows for evidence-based assignment of novel TSCs to existing gene annotations, and provides valuable information about overall transcript structure (Fig. 1B). We can thus attribute 82% of *D. melanogaster* TSCs to annotated genes, the remaining 18% potentially driving the expression of unannotated nonprotein-coding transcripts.

The comparison of biological replicates for the *D. melanogaster* time series confirms our ability to quantitatively measure promoter expression dynamics throughout development. Indeed, this analysis (Fig. S3) shows excellent reproducibility (Pearson R^2^ = 0.95) for TSCs with maximum expression >25 reads per million (RPM, Fig. 2A & S3), and very good reproducibility (Pearson R^2^ = 0.92) with maximum expression ≥10 RPM (Fig. S3).

**Figure 2:**
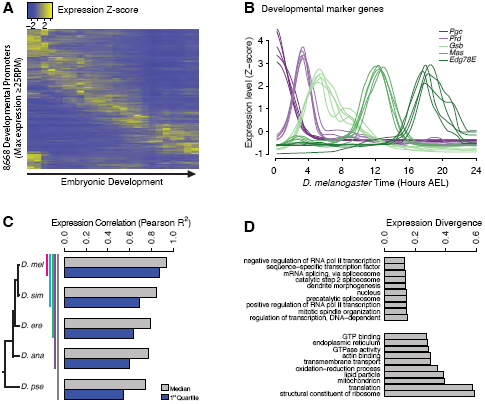
Evolutionary divergence of the developmental timing of promoter activity. (a) Hierarchically clustered expression profiles for individual *D. melanogaster* promoters throughout embryogenesis (24 time points; replicate 1). Only promoters with a maximum expression level >25 RPM (n=8,668), for which quantification reproducibility is very high, are included. (b) Expression profiles for 5 developmental marker genes, after global alignment of all time series to *D. melanogaster* (see Methods). The 5 curves for each gene correspond to the 5 species. (c) Conservation of the temporal expression profiles of individual promoters. For each subclade, we computed the average correlation coefficient between all pairs of species for each individual gene. The graph shows the median and first quartile over all genes with orthologs throughout the subclade. (d) The evolutionary divergence of expression specificity varies widely between Gene Ontology (GO) categories. For each gene with orthologs in all 5 species, maximum expression >25 RPM and expression changes >5-fold, we computed a measure of overall divergence across the clade (see Methods). The bar plot shows the average divergence by GO category, for the 20 categories with the lowest (top) and greatest (bottom) divergence.

Post-synchronization of developmental series was achieved by global alignments of all time series to the *D. melanogaster* reference to maximize the overall similarity between genome-wide expression profiles^42,43^. The resulting alignments for well-known developmental genes, used here as diagnostic markers, confirm the very high quality of the global alignments (Fig. 2B & S4). For all genes with one-to-one orthologs, the expression profiles are overall tightly conserved across species (Fig. 2C & S5A), but with substantial gene-to-gene variability: *hunchback,* for instance, displays considerable conservation in the expression of both of its promoters (Pearson R^2^ of 0.88 and 0.97; Fig. S5B), whereas the *RpL19* promoter shows rapid divergence (R^2^=0.08; Fig. S5C).

We found a strong relationship between selective constraints on expression specificity and gene function: indeed, the degree of expression divergence differs substantially between Gene Ontology (GO) annotation categories (Fig. 2D). Functions related to the regulation of transcription and splicing display the strongest conservation of expression, in accordance with the known molecular function of many master regulators of early development. Categories related to the core translational machinery and cytoskeletal structures display much more plastic expression specificities.

The high similarity of biological replicates, the accuracy of inter-species alignments for well-known developmental genes, and the biological features of evolutionary divergence patterns, together confirm our ability to accurately quantify promoter expression and its variation across species. Our observations also highlight the central importance of systems-level selective constraints, such as those acting on gene function and developmental stages, in shaping the evolution of gene expression.

### Core promoter structure defines broad developmental phases of gene expression

We leveraged our multispecies expression data to study promoter structure-function relationships, and thus gain insights into the regulatory code that determines developmental gene expression. We focused on a set of 3,462 promoters functionally active in all species that we classified either as housekeeping (<5-fold variation throughout development) or as developmentally regulated (≥60% of total expression within an 8-hour window). The developmentally regulated group was further clustered based on temporal correlation, yielding a total of 9 distinct expression clusters (Fig. 3A, see Methods). These thresholds were chosen to maximize the total number of promoters included and the similarity of profiles within each cluster, while still yielding clusters large enough for statistical analysis.

**Figure 3:**
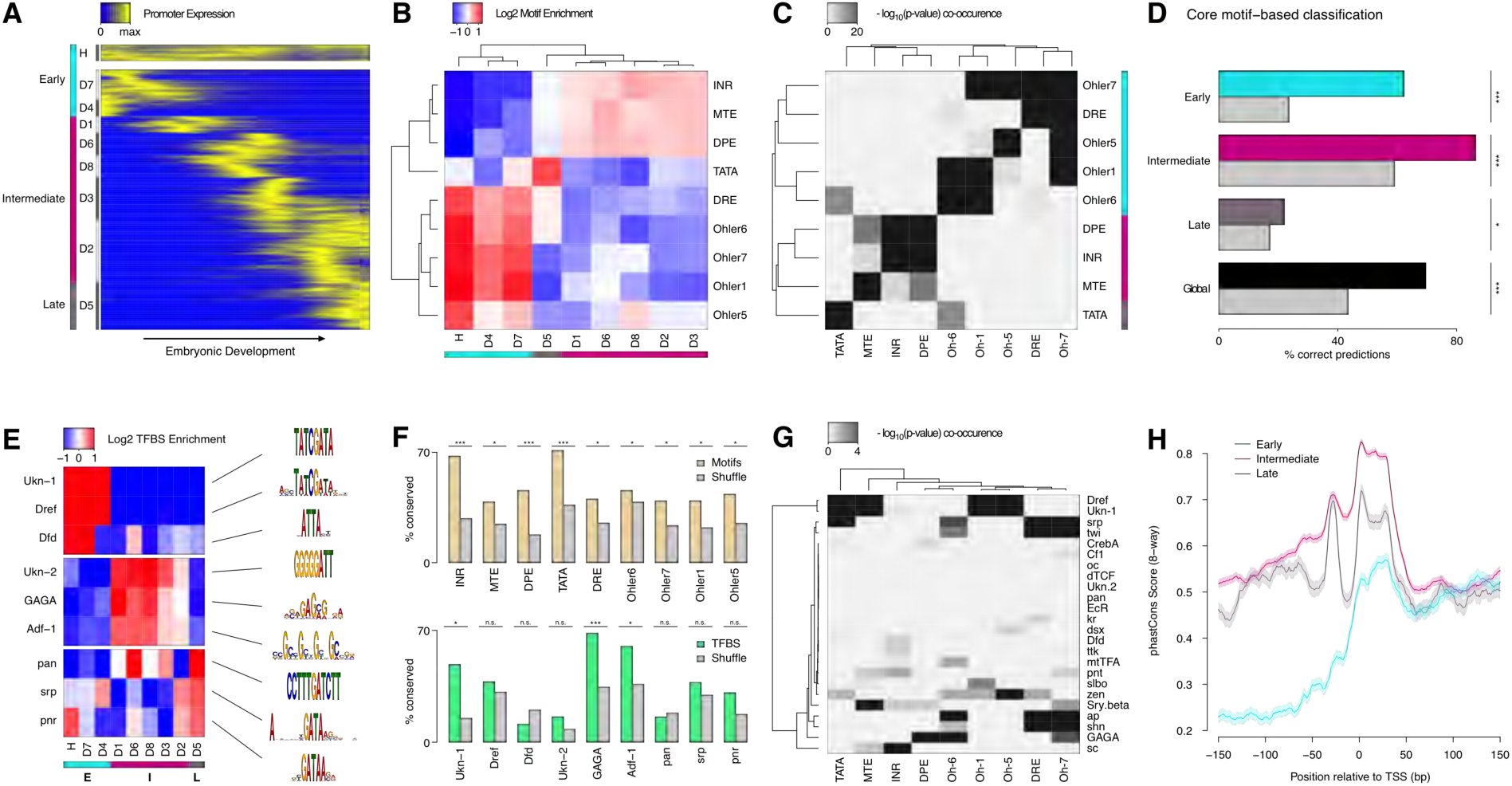
Core promoter structure defines 3 broad phases of embryonic development. (a) Clustering of expression profiles for 3,462 promoters (maximum expression >10RPM) defined as housekeeping (H) or developmentally regulated (D). Developmentally regulated promoters are divided into 8 groups based on hierarchical clustering of expression profiles (see Methods). Core promoter structure defines 3 broad developmental phases (color sidebar).^(b)^ Relative enrichment of core promoter motifs in each expression cluster. Three major clusters can be defined (bottom color bar), which correspond to 3 phases of embryonic development (see (a); Early: 817 TSCs; Intermediate: 2,047; Late: 598). (c) Clustering of core promoter motifs based on their co-occurrence in the same promoters. This approach yields the same 3 major sets of motifs previously defined based on expression profiles (see (b)). (d) Predictions of expression timing from core promoter structure. Quadratic discriminant analysis on log-transformed motif scores was used to predict the developmental phase in which promoters are expressed. The performance of the classifier in leave-one-out cross-validation (color bars) is compared to random expectation (grey bars; FDR-corrected chi-square tests applied to individual contingency tables as appropriate). (e) Distinct sets of TFBSs are enriched near the core promoters active in the 3 developmental phases. The top 3 motifs for each core promoter class are shown. (f) The conservation of core promoter motif composition between *D. melanogaster* and *D. pseudoobscura* confirms the biological relevance of the motifs. (Grey bars: conservation of shuffled motifs; FDR-corrected chi-square tests). (g) Many TFBSs are strongly associated with specific sets of core promoter motifs. Results are shown for the 3 motifs most enriched near the promoters of each expression cluster. (h) The 3 major core promoter classes display distinct profiles of sequence conservation. Lines: median phastCons scores across promoters of the class. Envelope: standard error in the estimate of the median, computed by bootstrapping.

These 9 expression clusters fall into 3 main groups, defined by the core motif composition of the promoters (Fig. 3B). Indeed, we unexpectedly observed robust enrichments for specific sets of motifs in the promoters of all individual expression clusters. Three major classes emerge, within which motif enrichments are relatively homogeneous (Fig. 3B).

Remarkably, these three classes define distinct temporal phases of embryonic development (Fig. 3A-B). The Early expression class, enriched for DRE and Ohler-1/5/6/7 motifs, consists of the promoters for maternally deposited transcripts, including the housekeeping cluster. The Intermediate class, enriched for Initiator (INR), MTE and DPE motifs, mediates transcription throughout a broad phase of mid-embryogenesis, from the onset of zygotic expression to the end of organogenesis. The Late class, enriched for TATA boxes, drives transcription around the transition to the first larval stage. Notably, expression clusters with vastly different specificities (e.g., D1 and D2) share the same promoter structure trends.

Importantly, clustering core motifs by their tendency to co-occur within the same promoters, regardless of expression timing, recapitulates the same three main motif groups (Fig. 3C). It does, however, also uncover a finer data structure, which suggests that there are a variety of promoter subtypes. In addition, we were able to train a classifier with substantial predictive power to distinguish the three promoter classes based solely on core motif scores (Fig. 3D). Taken together, our observations suggest that core promoter elements play a significant role in restricting windows of opportunity for expression during distinct periods of development.

Sequences proximal to the three core promoter classes are also enriched for different sets of TFBSs (Fig. 3E). The Early class preferentially harbors binding sites for Dref and Dfd, while the Intermediate class favors Trl (GAGA motif) and Adf-1. The Late class is enriched in pan, srp and pnr sites. The presence of most core promoter motifs and TFBSs is conserved between species far beyond random expectations (Fig. 3F), which validates the overall quality of our motif predictions. Interestingly, TFBSs are often specific for only a subset of expression clusters within a class (e.g., Dfd or GAGA). This suggests a model in which core promoter structure defines broad developmental periods of expression potential, and precise expression timing is then refined by sequence-specific transcription factors. Combinatorial encoding through stereotypical sets of TFBSs is likely to sharpen expression patterns even further (Fig. S6A-B).

The analysis of favored pairings between individual TFBSs and core promoter motifs suggests a possible mechanism to mediate this 2-step specification of expression patterns. Indeed, some TFBSs appear to be strongly associated with specific sets of core promoter motifs (Fig. 3G). Dref sites, for instance, are preferentially found along DRE and Ohler-1/6/7 core motifs. Likewise, twi and srp TFBSs, which often tend to be found together (Fig. S6A), have a robust association with INR and MTE. These strong affinities suggest that core motifs may tune the ability of a promoter to respond to specific sets of transcription factors. They may do so by recruiting different sets of general transcription factors (GTFs) that functionally interact with distinct groups of activators. Such a mechanism may channel various regulatory inputs to limited subsets of promoters and thus limit crosstalk between parallel pathways.

We found that the three promoter classes display markedly different profiles of sequence conservation (Fig. 3H & S6C). Importantly, this analysis only includes those promoters for which we have detected transcriptional activity in all 5 species, and we can therefore categorically rule out differences in rates of promoter gain/loss as an explanation. These observations suggest that the 3 promoter classes indeed have intrinsic structural differences that make them subject to distinct regimes of natural selection and sequence evolution.

### Selection on expression specificity shapes promoter sequence evolution

To further explore the selective pressures acting on regulatory sequences, we investigated quantitative relationships between the evolution of promoter structure (primary sequence) and function (expression specificity). We report a subtle but highly significant correlation between the conservation of promoter sequence and that of expression profiles (Fig. 4A-B). This shows that the effects of selection on expression specificity are reflected in the evolution of promoter sequences. The main areas of differential sequence conservation overlap regions preferentially occupied by precisely positioned core promoter elements (TATA, INR, DPE) and transcription factor binding sites (Fig. 4A). Importantly, this correlation does not hold for sequences >50bp downstream of the TSS, which are likely subject to additional selective pressures on 5’-UTR and protein-coding sequences. Promoters with highly divergent expression profiles are depleted of sites under purifying selection, and enriched for sites under positive selection (Fig. 4C-D).

**Figure 4:**
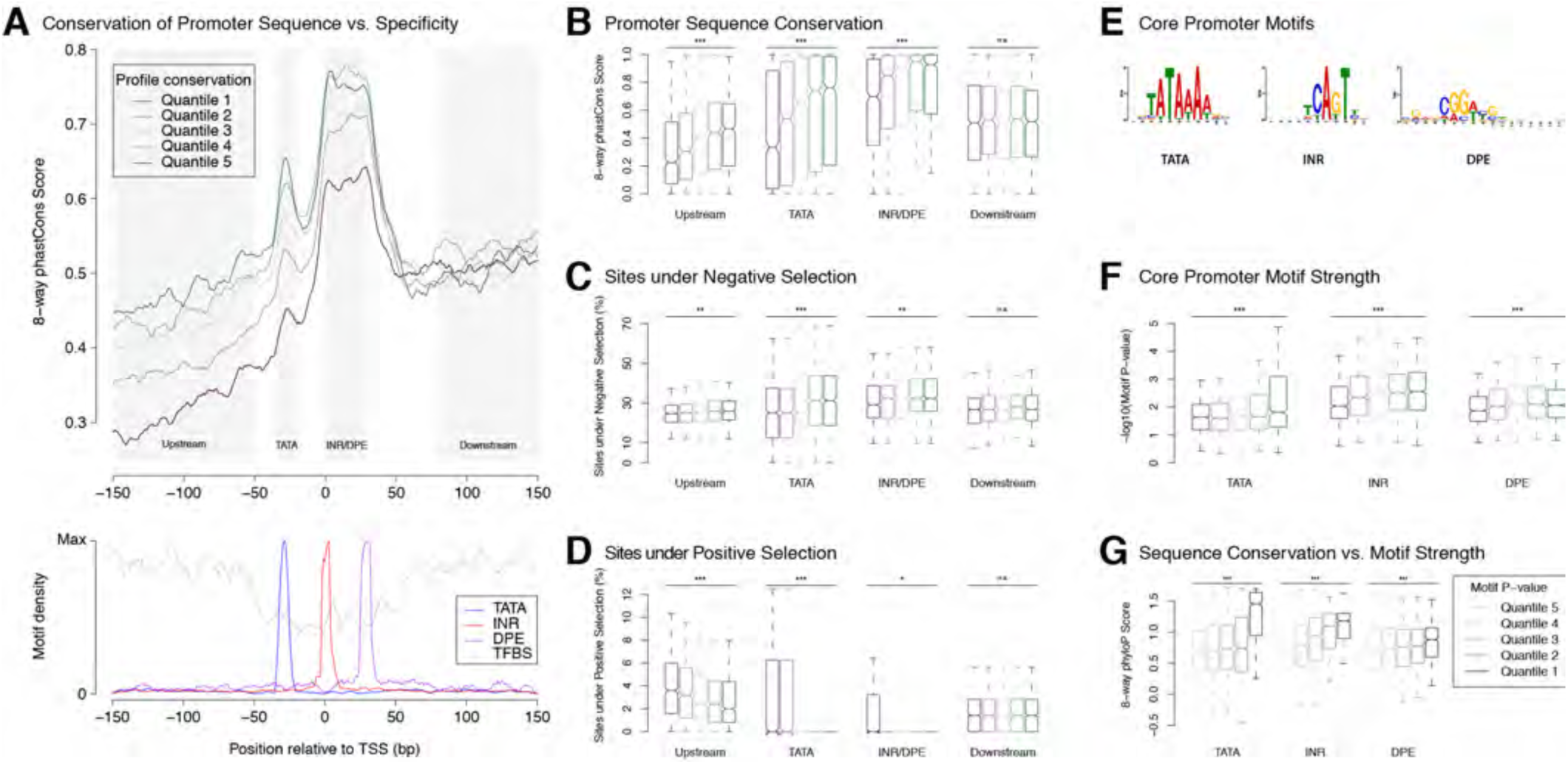
Selection on expression specificity shapes promoter sequence evolution. (a) Upper panel: average sequence conservation for *D. melanogaster* promoters, by expression profile conservation quantile. All *D. melanogaster* promoters with maximum expression ≥25RPM and functionally conserved in all 5 species were included (n=4,973). Lower panel: density of core promoter elements and TFBSs over all promoters. TATA box, INR, DPE: respective consensus sequences STATAAA, TCAGTY or TCATTCG, KCGGTTSK or CGGACGT^44^; TFBS motifs from Jaspar. (b) Sequence conservation by profile conservation quantile, over promoter subregions depicted as shaded areas in A. Upstream region runs from −300 to −50bp. (*** p-value < 10^-8^ for Pearson correlation test between profile conservation and sequence conservation; ** p < 10^-4^; * p < 10^-2^; *n.s.* not significant) (c) Proportion of bases under purifying selection (phyloP score >0.1). (d) Proportion of bases under positive selection (phyloP score <-0.1). (e) Core motif position weight matrices derived *de novo* from promoter sequences (see Methods). (f) Core motif strength correlates with expression profile conservation. (g) Core motif sequence conservation correlates with proximity to motif consensus.

*Ab initio* motif discovery within the regions of differential sequence conservation returned the canonical consensus sequences for TATA, INR and DPE (Fig. 4E). We found that promoters with highly conserved expression profiles tend to have core promoter elements whose sequence is closer to the motif consensus (Fig. 4F), and those in turn tend to be more conserved at the sequence level (Fig. 4G). Upstream flanking sequences also tend to show higher conservation of individual TFBSs (Fig. S7A). Importantly, it is possible to detect such a correlation for the binding sites of some individual transcription factors (Fig. S7B). This is a rather striking fact, considering that promoter-proximal enhancers generally bind more than one factor, and that many promoters are additionally regulated by distal enhancers not taken into account here. Finally, the conservation of promoter TFBS composition, as expected, also correlates with interspecific divergence (Fig. S7C).

These observations establish a clear relationship between core promoter sequence and expression specificity. Selective constraints on developmental expression timing act particularly strongly on core promoter elements, most notably Initiator, DPE and TATA elements. This is consistent with the idea that core promoter motifs are indeed key determinants of this specificity. Together, our results further support the hypothesis that core motifs and general transcription factors play a crucial role in determining promoter expression specificity.

### Promoter birth and death are widespread and dynamic

In addition to changes in the specificity of individual promoters, gene expression programs evolve through turnover of regulatory elements^36,45,46^. And indeed, we observed widespread birth and death of promoters throughout the clade: only 49% of *D. melanogaster* TSCs are functionally conserved in all 5 species (Fig 5A). To rule out genome assembly artifacts, we restricted our analysis to those with syntenic alignments to other genomes, and measured a functional conservation rate of 75% (Fig 5A). As some peaks lack alignments owing to genuine insertions or deletions, we expect the true conservation rate to be within the 49-75% range. Analyzing TSC conservation between species pairs, or from a *D. simulans-centric* perspective, yields similar conclusions (Fig. S8).

**Figure 5:**
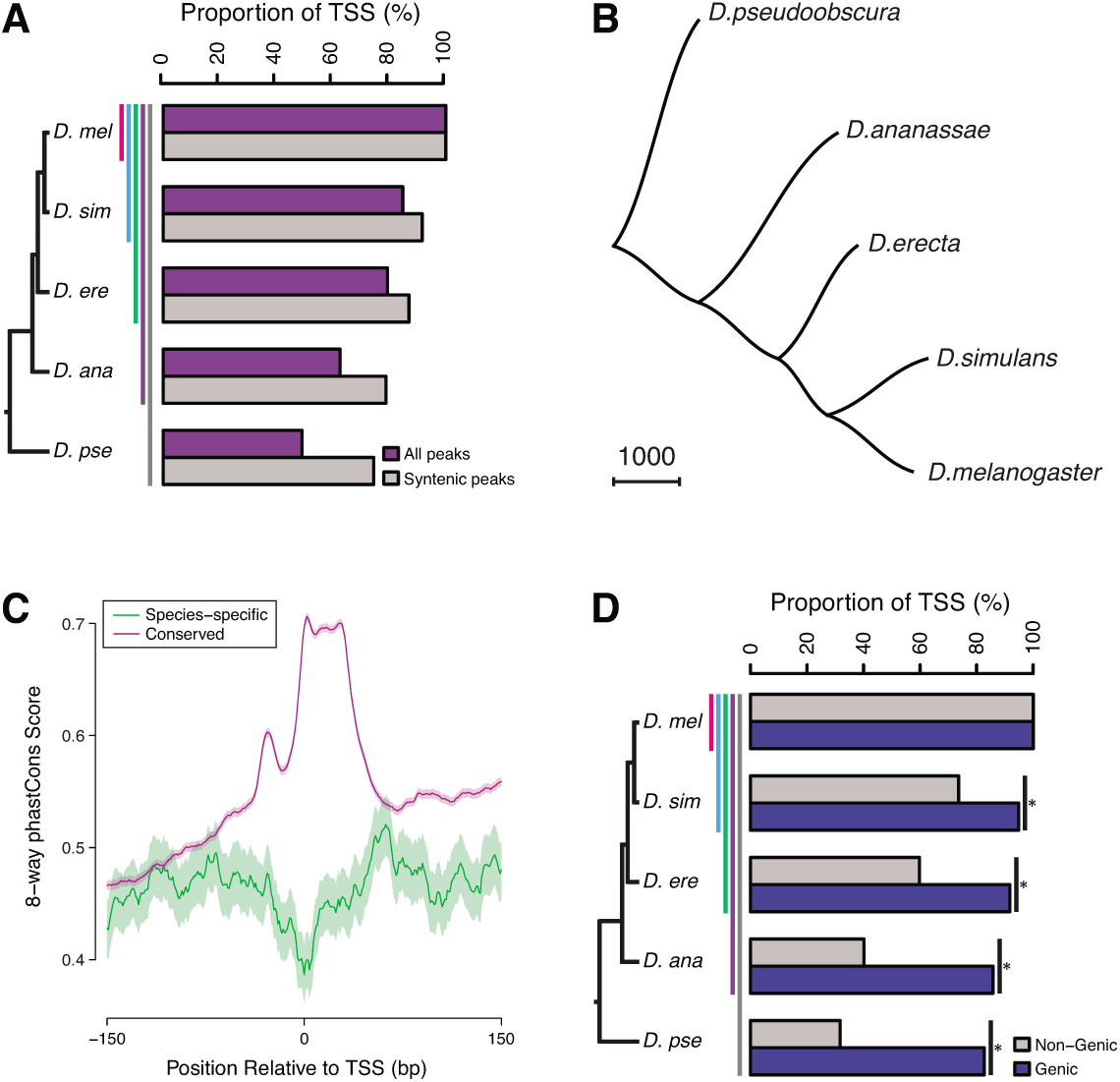
Widespread evolutionary gain and loss of promoters. (a) Proportion of *D. melanogaster* TSCs reproducibly detected in biological duplicates (first pair of bars) and functionally conserved in all species of subclades of increasing sizes. Subclades include all descendants of a common ancestor, and are designated by the species that is most distantly related to *D. melanogaster.* (b) The species phylogeny can be accurately reconstructed from patterns of TSC gain and loss. The presence/absence of each TSC was treated as a discrete character and the unrooted tree reconstructed using the Phylip software package. (c) Average profiles of sequence conservation over the TSCs functionally conserved across all 5 species and those specific to *D. melanogaster.* The shaded areas represent the standard deviation, estimated from 1,000 bootstraps. (e) TSCs driving the expression of annotated genes display a far higher degree of functional conservation than “orphan” TSCs (p<0.01 for all pairwise comparisons; chi-square test with Bonferroni correction).

The analysis of replicates for the *D. melanogaster* time series shows the false positive rate for gain/loss event detection to be under 0.1% (Fig. 5A). Although TSCs with lower expression levels tend to be less conserved, general trends are shared between TSCs of all expression levels (Fig. S9). Even though a gain/loss of expression during embryogenesis may reflect a shift in specificity rather than a complete gain/loss of function, the vast majority (91.4%) of *D. pseudoobscura* TSCs that were inferred to be lost in *D. melanogaster* based on embryo data are never expressed at any other stage of the life cycle (analysis of published data^37^). Therefore, we conclude that our strategy accurately and robustly detects promoter gain and loss events. Consistent with this, we can reconstruct the known species phylogeny by treating the presence or absence of individual TSCs as binary discrete characters in a standard parsimony framework (Fig. 5B). Functional conservation is reflected in sequence conservation: promoters active in all species display far higher sequence conservation than species-specific ones, which appear no more constrained than surrounding regions (Fig 5C & S10).

While purifying selection clearly plays a major role, a sizeable proportion of TSCs (25-50%) are not shared between all species, underscoring the inherently fluid nature of the regulatory landscape. Comparison to published data on the evolution of Twist binding sites^36^ revealed that overall, promoters evolve no more slowly, and possibly even faster, than TFBSs do (Fig. S11). Although a rigorous comparison between such disparate data types is delicate, this shows that promoters and TFBSs do not turn over on vastly different timescales.

Our data also reveals the existence of several thousand novel promoters that cannot be assigned to any annotated genes. Many of those may drive the expression of long noncoding transcripts, and it has long been a matter of debate to what extent this transcriptional activity plays meaningful biological roles. Hence we analyzed their rates of gain and loss to explore the selective pressures they may be subject to. We found a stark contrast in the degree of functional conservation of the two classes, with novel non-genic promoters evolving at a substantially higher rate than genic ones (Fig. 5D). There is, however, a very substantial proportion that is deeply conserved, suggesting the possibility of widespread functionality. The discrepancy between the classes may be due to a larger proportion of noncoding transcripts being devoid of biological roles and evolving neutrally. Alternatively, it may instead reflect a more pronounced tendency for noncoding transcription to take on lineage-specific roles and thereby be a driver of adaptation, as has been suggested before. To gain a better understanding of these questions, we sought to investigate in more depth the evolution of noncoding transcription.

### Deep conservation of long noncoding RNA promoters

The prevalence and biological relevance of noncoding transcription have long been major areas of contention. Attempts at resolving these issues using genomic sequence conservation have been largely inconclusive, probably due to minimal selective constraints on the primary sequence of these transcripts^27,28,47^ Studying promoter activity experimentally in a phylogenetic framework provides a unique opportunity to rigorously address the question of lncRNA conservation and functionality. Furthermore, our ability to pinpoint TSSs with single-nucleotide accuracy gives us unprecedented leverage to elicit elusive sequence conservation patterns. Furthermore, beyond sequence conservation, we are also in a position to assess selective constraints on the expression specificity of these promoters.

We found 3,682 embryonic TSCs in *D. melanogaster* that could not be functionally linked to any annotated protein-coding or small RNA gene, and could therefore represent putative lncRNA promoters. We also identified TSCs for 291 annotated lncRNAs, bringing the total up to 3,973. Their developmental expression kinetics appear to be diverse and exquisitely stage-specific (Fig. 6A).

**Figure 6:**
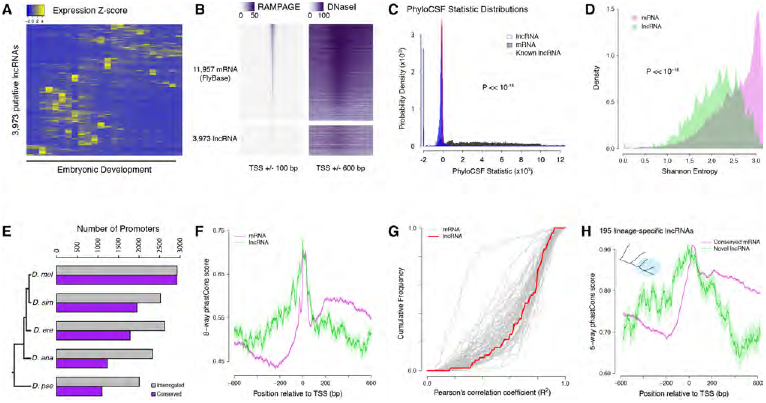
Strong purifying selection on long noncoding RNA promoters. (a) Developmental expression profiles of putative lncRNA promoters (n=3,973). (b) Heatmaps of RAMPAGE signal (left; number of reads) and DNase-seq signal (right; arbitrary uits) over individual TSCs. We are comparing TSCs that overlap FlyBase-annotated mRNA transcription start sites (top), which we use as positive controls, to the TSCs of putative lncRNAs (bottom). In each group, TSCs are sorted by total RAMPAGE signal intensity. (c) Distribution of phyloCSF scores for transcript models corresponding to putative lncRNAs (n=1,475) and mRNAs (n=16,105). Transcript models were built from publicly available RNA-seq data using Cufflinks. The phyloCSF metric quantifies the protein-coding potential of transcripts, based on the presence and conservation of ORFs. (d) Shannon entropy of the temporal expression profiles for lncRNA (n=2,397) and mRNA (n=18,067) promoters with maximum expression ≥ 2RPM. Overall, the profiles of lncRNA promoters have lower entropy, reflecting more acutely stage-specific expression. (e) Number of *D. melanogaster* lncRNA promoters functionally preserved in other species. The grey bars represent the number of promoters for which the multiple sequence alignments passed our filtering criteria, and therefore could be interrogated. (f) Promoter sequence conservation. Considering all promoters that are functionally preserved in all 5 species, the sequences of lncRNA promoters (n=631) are under comparable selective pressure to those of protein-coding genes. (g) The developmental expression profiles of functionally conserved lncRNA promoters are far more constrained than those of many categories of protein-coding genes. We considered all promoters with maximum expression ≥ 25 RPM and expression changes ≥ 5-fold (n=55 lncRNA promoters). (h) We identified 195 *D. melanogaster* lncRNA promoters that are functionally preserved within the *melanogaster* subgroup, but not in the 2 outgroup species, and are therefore likely to have been recently acquired specifically in this lineage (inset, top left). Lineage-specific lncRNA promoters display a level of sequence conservation within the subgroup similar to that of conserved protein-coding gene promoters.

The analysis of published genome-wide DNaseI hypersensitivity data^48^ confirmed that these putative lncRNA TSCs are likely to correspond to genuine promoters (Fig. 6B & S12). In addition, to verify that the transcripts expressed from these TSCs are indeed independent and devoid of any significant protein-coding potential, we built transcript models from a recently published RNA-seq developmental time course^25^. We successfully generated transcript models for 16,105 TSCs, including 1475 lncRNA TSCs. Most of them appear to correspond to full-length transcripts, and the vast majority of putative lncRNAs do not overlap annotated protein-coding sequences (Fig. S13). Analyses of protein-coding potential confirm that the overwhelming majority of transcripts are unlikely to encode proteins, or even peptides as short as 10 amino acids (Fig. 6C & S13). Transcripts from 18 loci are likely to encode short open reading frames (sORFs, <100 residues). We conclude that the vast majority of candidate transcripts are likely to be genuine lncRNAs.

The expression profiles of lncRNA promoters have unique properties that set them apart. As a class, they tend to be substantially more stage-specific than their protein-coding gene counterparts, as measured by the Shannon entropy of their activity profiles (Fig. 6D). Comparing their developmental expression timing to that of protein-coding gene promoters suggests a broad diversity of potential developmental functions (Fig. S14 & Table S1).

The detection of lncRNA TSCs is highly reproducible across *D. melanogaster* biological replicates (Fig. 6E). Of all *D. melanogaster* lncRNA TSCs, 2,016 can be aligned to the *D. pseudoobscura* genome assembly and 1,111 are functionally conserved (Fig. 6E), suggesting that they have been maintained since the last common ancestor of these two species. In order to investigate whether lncRNA promoters are under purifying selection, we focused on an extremely stringently selected set of 631 TSCs that are active in all 5 species. This set includes well-known essential noncoding transcription units, such as *bithoraxoid^9,50^* (Fig. S15) and *roXl*^51^. Overall, the level of sequence conservation at these functionally preserved lncRNA promoters is similar to that observed at protein-coding gene promoters (Fig. 6F). Their developmental expression specificity is also more constrained than that of many protein-coding gene promoters (Fig. 6G). Both observations taken together argue strongly for sustained selective pressure on these 631 lncRNA promoters for 25-50 million years. Furthermore, many TSCs of interest were excluded from this analysis simply because of the poor quality of genome assemblies, and this is therefore an extremely conservative set. Therefore, to place a more reasonable lower bound on the true number of conserved lncRNA promoters, we focused on those shared between the 3 species of the *melanogaster* subgroup. These 1,529 promoters similarly display a high degree of sequence conservation within the subgroup, and their expression specificity also appears constrained (Fig. S16).

Still, it remains that lncRNA promoters as a class evolve much faster than those of protein-coding genes (Fig. 5D), and it has been a matter of debate whether this reflects lineage-specific functions or merely neutral evolution. To address this question, we focused on 195 lncRNA TSCs that are specific to the *melanogaster* subgroup, despite the orthologous sequences being present in the genomes of the two outgroups (Fig. 6H). Surprisingly, they display the same degree of sequence conservation as the protein-coding gene promoters that are shared throughout the subgroup (Fig. 6H). The assessment of conservation at orthologous sequences in outgroup species confirms that the selective constraints are indeed lineage-specific (Fig. S17). This argues that these evolutionarily recent lncRNAs have come under purifying selection after acquiring lineage-specific functions.

Taken together, our observations show that a vast proportion of lncRNAs are indeed under purifying selection for biological functions relevant to embryonic development.

### *schnurri-like RNA:* A deeply conserved, developmentally regulated lncRNA gene

To validate our findings in one specific case, we focused on the *FBgn0264479* locus, which displays one the most tightly conserved expression patterns among all the lncRNA genes in our dataset. This is an intriguing embryonic transcript that, although it has been annotated based on expressed sequence tag (EST) data, has to our knowledge never been characterized. The 0.5kb FBgn0264479 RNA is extremely unlikely to encode functional peptides, as assessed by phyloCSF analysis (score of −217.3) and manual curation (Fig. S18). It is highly expressed in all 5 species surveyed, in a strikingly conserved temporal pattern restricted to a ∼3-hour period encompassing the onset of gastrulation (Fig. 7A-B & S19). Northern-blot analysis confirmed the size and expression dynamics of the transcript (Fig. 7C & S20)

**Figure 7:**
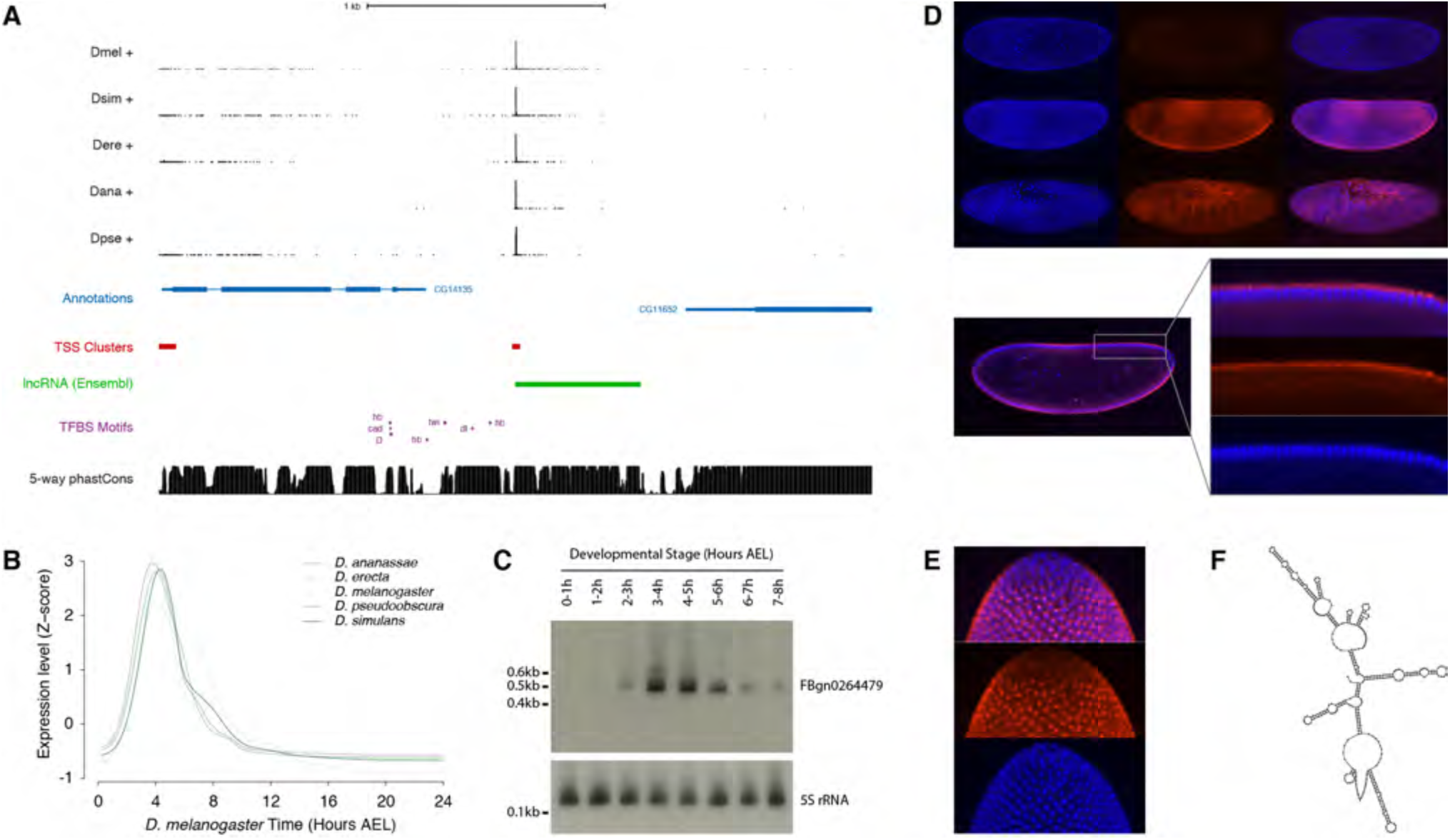
Functional characterization of *schnurri-like RNA.* (a) The *schnurri-like RNA* locus (*FBgn0264479*; UCSC Genome Browser). Data tracks, from top to bottom: RAMPAGE signal in 5 species (5 upper tracks), FlyBase gene annotations (blue), *D. melanogaster* TSCs (red), transcript annotation from Ensembl (green), TFBS motif predictions around the *slr* promoter (purple), sequence conservation within the *melanogaster* subgroup (black). (b) slr transcript expression profiles in 5 species. (c) Northern-blot against the slr RNA. The 5S ribosomal RNA was used as a loading control (lower panel). (d) RNA-FISH for the slr transcript. Upper panel, top to bottom: maximum-intensity projections of confocal series for embryos at stages 4, 5 and 7 (Blue: DAPI, red: FISH). Bottom panel: Single confocal section from the embryo in the middle of the top panel. Controls with sense probes showed very little background. (e) slr RNA-FISH; lateral view of the posterior pole of a stage 5 embryo. (f) Mfold-predicted secondary structure of the RNA.

The body of the transcription unit displays hallmarks of robust purifying selection within the *melanogaster* subgroup (Fig. 7A). In addition, publicly available chromatin immunoprecipitation data^52,53^ reveals the binding of several transcription factors to the promoter region (Fig. S19), and their putative binding sites identified by sequence motif search also show evidence of purifying selection (Fig. S21). The expression dynamics of the transcription factors are consistent with their regulating the *FBgn0264479* promoter (Fig. S21).

Fluorescent *in situ* hybridization (FISH) in early embryos revealed expression along the ventral and dorsal midlines at the late blastoderm stage, with the exclusion of lateral regions, the primordial germ cells and the prospective head (Fig. 7D). Based on this distinctive expression pattern, highly reminiscent of the well-characterized *schnurri* gene^54^, we renamed this lncRNA gene *schnurri-like RNA* (*slr).* This early expression domain subsequently evolves into a complex segmented pattern by the end of germband extension (Fig. 7D). In the late blastoderm, the RNA is found almost exclusively in the cytoplasm at the apical pole of the cells (Fig. 7D). It appears at that stage to be generally concentrated in a single major focus per cell (Fig. 7E). Secondary structure predictions show that the slr transcript is likely to be highly structured (Fig. 7F), suggesting a high potential for RNA-protein interactions.

Although further characterization will be required to decipher the precise biological roles of this lncRNA, our observations confirm its noncoding nature, definitively establish its developmental expression specificity, and provide clear evidence that it has been under strong purifying selection over at least 25-50 million years.

## Discussion

This work provides, to our knowledge, the first genome-wide overview of promoter evolution in *Drosophila*, and we leveraged this comparative framework to study the sequence determinants of developmental expression specificity. Through nucleotide-resolution mapping of TSSs and quantitative measurements of expression kinetics, our study yields new insights into the transcriptional regulatory code. We find that distinct classes of core promoters drive transcription in three broad phases of embryonic development. Each class is defined by a characteristic set of core motifs, and is associated with regulation by specific groups of transcription factors. Of note, we successfully detected *in vivo* some functional associations between Dref and housekeeping promoters, and between Trl and some developmentally regulated promoters, that were recently described *in vitro*^18^ (Fig. 3). Our analysis generalizes the concept of specific interactions between core promoters and enhancer sequences, and demonstrates for the first time its global relevance in a developmental context.

We propose a hierarchical model for transcriptional regulation in which core promoter syntax defines broad temporal windows of opportunity for activation, and precise expression timing is subsequently refined by the binding of sequence-specific activators and repressors at enhancers. Core promoters may restrict regulatory inputs by recruiting different sets of general transcription factors (GTFs) that functionally interact with distinct groups of transcription factors^20^. Some GTFs have been shown to shape the expression specificity of individual promoters, and it is known that different activators and repressors have distinct requirements for cofactors and GTFs^18,20^. Such a mechanism may channel regulatory inputs to limited subsets of promoters, and thus limit crosstalk between promoters and enhancers across the genome. Notably, what applies to time may also apply to space, and it is possible that similar core promoter/enhancer interplay hierarchically specifies gene expression in broad developmental lineages and individual cell types.

Evolutionary analysis of developmental expression specificity further supports this model. First, the three major classes of core promoters defined here show drastically different patterns of sequence evolution, suggesting substantial differences in their underlying structure and functional interactions with the transcriptional machinery. At a finer scale, the conservation of expression specificity across species correlates with the degree of sequence conservation at canonical core promoter elements, such as TATA boxes and Initiator motifs. This is highly suggestive of an instructive role for these sequence elements, once thought generic, in defining developmental gene expression patterns.

Approximately 4,000 promoters were found to drive the expression of lncRNAs during embryogenesis, a strikingly high number for this very brief developmental period. Our findings likely apply to other developmental stages as well, as we previously reported the existence of 7,421 putative lncRNA promoters in an analysis of the whole *D. melanogaster* life cycle^37^. In addition, we detected the expression of only 205 of 1,119 recently identified lncRNAs^27^. This suggests that we are only beginning to scratch the surface of lncRNA biology in *Drosophila.* Importantly, we show here that vast numbers of these promoters are under strong selective pressure, at the levels of both promoter sequence and expression specificity. A *melanogaster* subgroup core set of at least 1,529 is under substantial selective constraint, and most of those are therefore highly likely to have biologically relevant activities.

In agreement with previous reports ^28,47,55^, we find that lncRNA genes evolve faster than their protein-coding counterparts. It has been an unresolved debate so far whether this reflects neutral evolution or lineage-specific functions. Our observation that lineage-specific lncRNAs are also under substantial selective pressure reveals that noncoding transcription may be a major driver of phenotypic diversification and organismal adaptation. We recently showed that transposable elements play an important role in the evolutionary gain of promoters, and in particular of lncRNA promoters^37^. We propose that transposon proliferation is a major mechanism favoring the neofunctionalization of intergenic regions as sources of biologically active noncoding transcripts.

To open a window into the biology of developmentally regulated lncRNAs, we focused on *schnurri-like RNA,* a deeply conserved yet never-before characterized gene. We experimentally validate the existence and expression kinetics of the slr transcript, and demonstrate that both its promoter sequence and its developmental expression specificity are deeply conserved across drosophilids. Interestingly, this lncRNA is expressed in a spatial profile highly similar to that of the *schnurri* gene, which is part of the Dpp (TGF-β/Smad) signaling pathway and plays an essential role in the establishment of dorsoventral embryo polarity^54,56^. The punctate cytoplasmic localization pattern of the slr lncRNA is reminiscent of the targeting of multiple Dpp pathway components to endosomes in larval wing discs^57-59^. Endosomes localize apically in the late embryonic blastoderm^60^, and mutants for *Sara*, the Smad endosome-targeting factor^57,59^, die early in embryogenesis^57^, suggesting that endosome-based signaling is also essential at that stage. Taken together, our observations are suggestive of a possible role for the slr lncRNA in TGF-β signaling, a crucial pathway in all animals that plays a major role in human disease, including cancer.

In recent years, it has become clear that noncoding transcription serves a myriad of molecular functions in Eukaryotes, and plays a part in virtually every known biological process^29-32^. LncRNAs have been shown to regulate transcription and chromatin structure, as well as mRNA stability and protein localization. Sometimes it is the transcription of the locus itself that plays a mechanistic role, rather than the resulting transcript - as in the case of the upstream *bxd* promoter^50^, which has one of the most highly conserved expression profiles that we have observed. Our work unambiguously demonstrates the biological relevance of noncoding transcription to developmental processes, and establishes *D. melanogaster* as an excellent model for exploring the diverse functions of lncRNA genes. We expect that systematic efforts on a larger scale will illuminate the biology of a long-ignored class of genes that has proven its worth.

## Author Contributions

P.J.B and T.R.G. conceived the project and designed experiments. P.J.B carried out experiments and data analysis. P.J.B and T.R.G. wrote the manuscript.

## Acknowledgements

The authors would like to thank Alexander Dobin, Felix Schlesinger, Chris Zaleski, Carrie Davis and all other members of the Gingeras group at CSHL for their assistance and advice, as well as Richard McCombie and the CSHL sequencing facility for their services. We also thank Alexander Gann, Gregory Hannon, Zachary Lippman, Joshua Dubnau, Adrian Krainer, Brenton Graveley and Mike Levine for helpful discussions and advice. We are grateful to Thomas Kaufman and the FlyBase team for their permission to reproduce images. Work supported in part by the National Human Genome Research Institute, modENCODE Project, contract U01HG004271.

## Data Availability

The primary data for this study is available through the GEO database, under accession numbers GSE36212 and GSE89335. GSE36212 is public, and reviewers can access GSE89335 here: https://www.ncbi.nlm.nih.gov/geo/query/acc.cgi?token=onklcqcedfohvkt&acc=GSE89335

## Online Methods

### Fly stocks & Embryo collections

All Drosophila strains were obtained from the Drosophila Species Stock Center at UC San Diego, CA (https://stockcenter.ucsd.edu/info/welcome.php). For each species considered we worked with the reference genome strain. Stock numbers: *D. melanogaster* #14021-0231.36, *D. simulans* #14021-0251.195, *D. erecta* #14021-0224.01, *D. ananassae* #14024-0371.13, *D. pseudoobscura* #14011-0121.94. Stocks were maintained on standard cornmeal medium. Embryo collections were performed in population cages (Flystuff, #59-116). 2- to 7-day-old flies were left to acclimatize to the cage for at least 48h and regularly fed with grape juice-agar plates (Flystuff, #47-102) generously loaded with yeast paste. After two 2-hour pre-lays, embryos were collected in 1-hour windows and aged appropriately (24 time points, 0-24h). Embryos were washed with deionized water, dechorionated for 90 sec with 50% bleach, rinsed abundantly with water, and snap-frozen in liquid nitrogen.

### RNA Extraction & RAMPAGE Library preparation

Total RNA was extracted from embryos using a Beadbeater (Biospec, Cat. #607) with 1.0 mm zirconia beads (Biospec, #11079110zx) and the RNAdvance Tissue kit (Agencourt #A32649) according to the manufacturer’s instructions, including DNaseI treatment. We systematically checked on a Bioanalyzer RNA Nano chip (Agilent) that the RNA was of very high quality. Libraries were prepared as described before^37,41^. 5’-monophosphate transcripts were depleted by TEX digest (Epicentre #TER51020). For every time series, each sample was labeled with a different sequence barcode during reverse-transcription, and all samples for the series were then pooled and processed together as a single library. Quality control and library quantification were carried out on a Bioanalyzer DNA High Sensitivity chip. Each library was sequenced on one lane of an Illumina HiSeq 2000.

### Genome references & annotations

All reference sequences and annotations were obtained from Flybase (http://flybase.org). *D. melanogaster* release 5.49, *D. simulans* r1.4, *D. erecta* r1.3, *D. ananassae* r1.3, *D. pseudoobscura* r2.9.

### Primary data processing

Reads were mapped to the appropriate reference genomes using the STAR aligner^61^. Peaks were called on the pooled data from whole time series, using a custom peak-caller described previously^37,41^. We used parameters optimized to yield good TSS specificity with respect to annotations and comparable numbers of peaks for all species. All peaks overlapping FlyBase-annotated rDNA repeats were filtered out.

### TSC conservation

Functional conservation was assessed for all peaks with ≥15 RAMPAGE tags that did not map to heterochromatic regions or chr4 in *D. melanogaster,* or orthologous regions in other species. We translated the genomic coordinates of each peak in each species to coordinates in the multiple sequence alignment of all genomes (15-way MultiZ alignment from UCSC, http://hgdownload.soe.ucsc.edu/goldenPath/dm3/multiz15way). To be considered for analysis, each peak was required to have a unique syntenic alignment in all other species considered, defined as follows: both ends of an 800-bp window centered on the middle of the peak had to map to the same strand of the same chromosome or scaffold, 50% of bases had to be aligned *(i.e.,* not in assembly gaps), and 25% of bases had to align to orthologous bases (not alignment gaps). Raw 5’ signal for each genome was also translated into multiple alignment coordinates. For each peak from each species, functional conservation was assessed by counting the number of RAMPAGE tags in each species. A peak was considered absent in a target species if it had at least a 100-fold lower signal than in the reference species. Peaks with <100 tags in the reference species were considered absent if they had no detectable signal in a target species.

### Phylogeny reconstruction

The peaks from all species were merged and collapsed in multiple alignment space to generate a nonredundant set of all peaks in the clade. The conservation of these peaks was assessed as described above. The phylogenetic tree was inferred by treating the presence/absence of each peak as a 2-state discrete character, sequentially using the MIX and PARS program of the PHYLIP suite according to the recommendations of the software documentation (http://evolution.genetics.washington.edu/phylip.html).

### Sequence conservation

Per-base conservation scores were computed by running the phastCons and phyloP programs of the PHAST suite v1.1 on the MultiZ alignment according to the recommendations of the software documentation (http://compgen.bscb.cornell.edu/phast). Depending on the subclade of interest, some species were excluded from the alignment for certain analyses. Pre-computed phastCons scores for the full 15-way alignment were downloaded from UCSC (http://hgdownload.soe.ucsc.edu/goldenPath/dm3/phastCons15way).

### Core promoter motifs

For analyses of motif composition, we only considered *D. melanogaster* TSCs that were functionally conserved across all 5 species. We used pairwise chained alignments downloaded from UCSC (http://hgdownload.soe.ucsc.edu/downloads.htmltffruitfly) to align the most heavily used position of each TSC *(i.e.,* the main TSS) to all other genomes. Peaks for which the maximum position could not be aligned to all genomes were excluded from the analysis. A custom script was used to search for matches to previously characterized core promoter motifs^44^ within a 301-bp window centered on the main TSS. Consensus sequences for sets of peaks with matches to individual motifs were computed using MEME v4.9.0 (http://meme.nbcr.net/meme).

### Time series alignment

Z-score transformed gene expression time series from all species were registered to one another using the GTEM suite ^42^ according to the recommendations of the software documentation (http://flydev.berkelev.edu/cgi-bin/GTEM/index.htm). One-to-one orthology calls from Flybase (2012 release 2) were used to match gene expression profiles between species. We pre-processed pairs of datasets *(D. melanogaster* and another species) to compensate for differences in annotation quality and peak calling between species. We identified orthologs of TSCs that had detectable expression (≥10 tags) but initially failed to be called in one species. In addition, when a functionally conserved TSC had been attributed to an annotated gene in one species but not the other, we corrected this discrepancy by attributing it to the gene in both species. For the *D. ananassae* dataset, the 8^th^ time point failed to yield acceptable data, and was excluded from the analysis. All time series were upsampled 5-fold and smoothed with a 2-hour window size using RZ-Smooth v4.1. Optimal global alignment paths between *D. melanogaster* and the other datasets were computed with T-Warp v3.2 with Pearson distance matrices (3-hour window). M-Align v2.8 was used to align each series to the *D. melanogaster* reference and smooth the final aligned series (1-hour window). The expression profiles of individual TSCs were registered to one another with M-Align, using the optimal alignment path computed for gene expression profiles. Prior to alignment, we used the UCSC liftOver tool

(http://hgdownload.cse.ucsc.edu/admin/exe/linux.x8664/liftOver) to identify *D. melanogaster* TSCs that aligned well (≥50% of bases aligned) to all other genomes. The temporal expression profiles of those orthologous genomic positions only were aligned.

### Expression profile conservation

We defined the conservation of individual expression profiles (TSCs or genes) across a clade as the average Pearson R^2^ for all pairwise comparisons of species within the clade.

### Clustering of TSC expression profiles

We classified all *D. melanogaster* TSCs with maximum expression ≥10RPM (n=11,900) as either Housekeeping (<5-fold variation throughout the time series, n=587) or Developmentally Regulated (≥60% of total expression within an 8-hour window, n=6,015). We further selected TSCs functionally conserved in all 5 species (n=240 and n=3,824, respectively). Developmentally regulated TSC profiles were hierarchically clustered (R *hclust,* distance metric *1 - cor(t(expr), method="pearson"))* and initially grouped into 12 clusters (R *cutree,* k=12). After filtering out excessively small clusters (<200 TSCs, n=4), further analysis was conducted on the remaining 8 regulated clusters (n=3,222 TSCs). See Fig. 3A for clustering results.

### lncRNA transcript reconstruction & phyloCSF ORF analysis

We ran Cufflinks (v2.2.1) independently on each dataset of a published RNA-seq developmental series. Cuffmerge was used to generate a consensus annotation set. Transcript models were attributed to a RAMPAGE TSC if their 5’ end lay within 150 of that TSC. Models without a matching TSC were excluded from further analyses. phyloCSF was run on these annotation sets and the 15-way multiZ whole-genome alignments.

### Analysis of sequence motifs

We used the MEME Suite v4.9.0 (primarily FIMO and MAST) to search promoter regions for a previously published compendium of motifs^62^ corresponding to core promoter motifs and TFBSs. MEME was used to generate consensus motifs from specific subregions of RAMPAGE-defined promoters (Fig. 4).

### Other software

Custom analysis scripts were written in Python 2.7 (http://www.python.org). R was used for graphics generation (http://www.r-project.org).

### *FBgn0264479* promoter sequence analysis

We identified potential regulators of the FBgn0264479 promoter based on the BDTNP transcription factor ChIP-chip data available from the UCSC genome browser website^52,53^. For 10 factors that bind the promoter in embryos (bcd, cad, D, dl, ftz, gt, h, hb, Kr, twi), we searched the 600bp upstream of the main TSS for Jaspar TFBS motifs (http://jaspar.genereg.net), using FIMO. We identified 7 motifs at a p-value cutoff of 2x10^-4^.

### Northern-blot

We ran 8μg of total RNA per sample on an 8% acrylamide 8M urea gel with a Invitrogen Novex minigel system. Transfer to a nylon membrane was carried out in 0.5X TBE in a Novex XCell II module, followed by UV-crosslinking (1,200J). For detection, we used a combination of 6 oligonucleotide probes targeting FBgn0264479 (5’-gaacatcgcttgcagtgcag, 5’-cgatggatgttgtcggtcgg, 5’-ctctcgttctttgattcttc, 5’-caggatgtgtggtgttccac, 5’-agattggatccttatggttg, 5’-atatgctgacactgcatggt). 30 pmol of oligo mix were radioactively labeled with γ^32^-ATP and PNK. Following phenol-chloroform extraction, the labeled probes were hybridized for 2 hours at 42°C in 40mL of ULTRAHyb buffer (ThermoFischer). After serial washes with decreasing concentrations of SSC buffer (final stringency 0.5X), the membrane was exposed on Kodak BioMax autoradiography film. After stripping and control re-exposure, a similar protocol was used to detect the 5S rRNA on the same membrane, using a single probe (5’-caacacgcggtgttcccaagccg).

### Fluorescent *in situ* hybridization

Templates for probe synthesis were generated by amplification of FBgn0264479 cDNAs with primers 5’-CGATGTTCTCCGACCGACAA and 5’-TGCACTACTTAGACTAAATTGGCT. In separate reactions, a T3 promoter sequence was added at either one end or the other, for the generation of sense and antisense probes. Amplicons were cloned into a TOPO-T/A vector (Life Technologies #K4575-01) and checked by Sanger sequencing. RNA-FISH was performed on 0-5 hours AEL *y; cn b sp* embryos as described before^63^, and imaged on a Perkin-Elmer UltraVIEW VoX confocal microscope. Biotin-conjugated mouse monoclonal anti-DIG (Jackson ImmunoResearch Laboratories Inc., Cat. No. 200-062-156) previously validated for this application^63^.

### Sample size estimates

In this work, relevant comparisons are between groups of promoters or groups of genes. All comparisons were designed to include all TSCs or genes of interest throughout the genome (e.g., all lncRNA TSCs *vs.* all genic TSCs), while applying expression level thresholds calibrated on the analysis of *D. melanogaster* replicates to ensure measurement reproducibility.

### Code availability

RAMPAGE analysis pipeline and custom analysis scripts (Python & R) available upon request.

---

**Figure 1.**
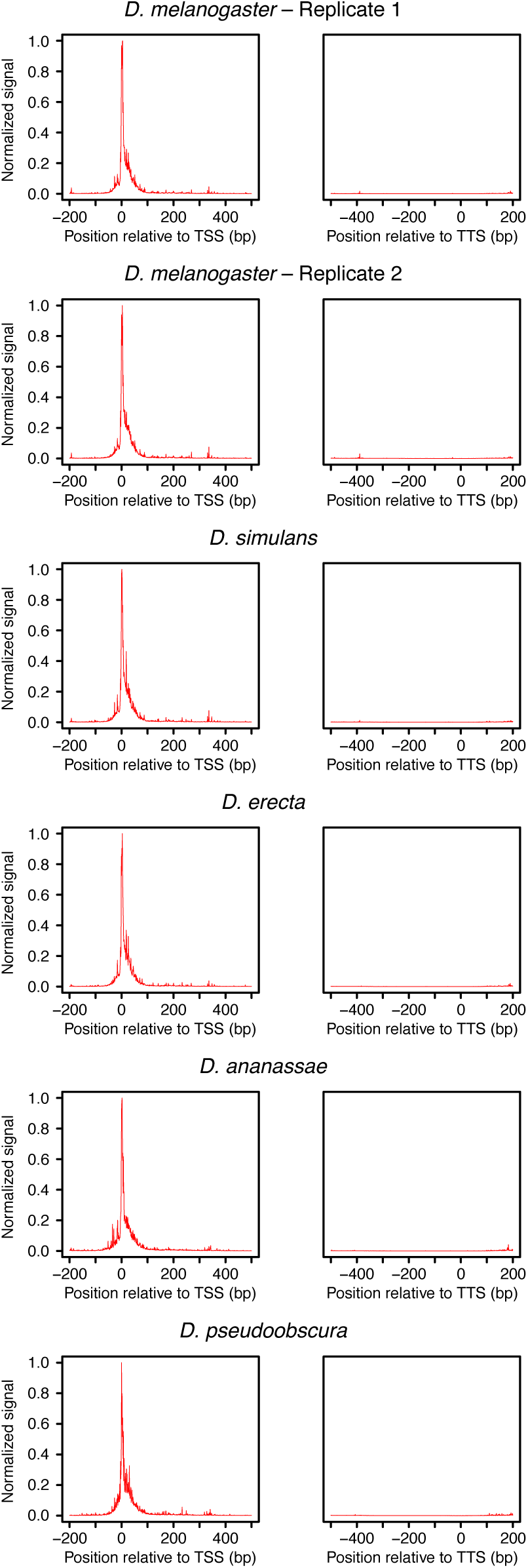
Distribution or raw RAMPAGE signal over transcript annotations. For each species, RAMPAGE reads were mapped to the appropriate genome. The raw 5’ signal was then converted to orthologous *D. melanogaster* coordinates using chained pairwise alignments from UCSC. Metaprofiles were constructed by summing signal intensity over FlyBase r5.49 mRNA annotations.

**Figure 2.**
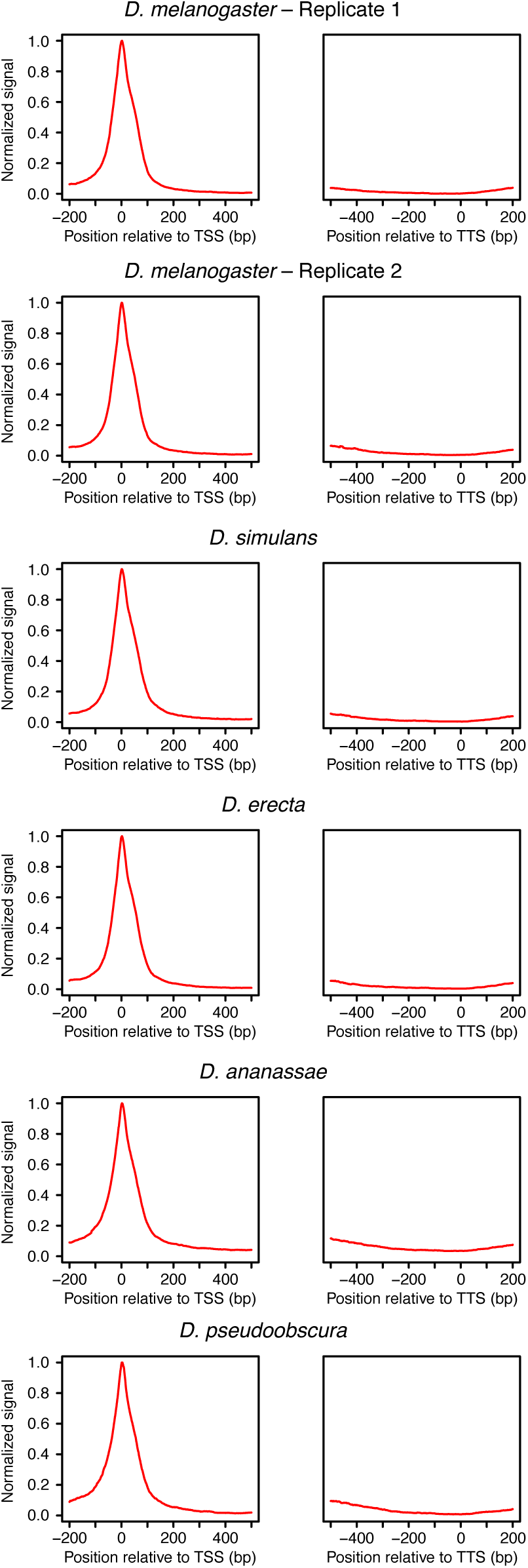
Distribution or RAMPAGE peaks over transcript annotations. For each species, RAMPAGE reads were mapped to the appropriate genome and peaks called as described in Methods. The peak coordinates were then converted to orthologous *D. melanogaster* coordinates using chained pairwise alignments and the liftOver tool from the UCSC Genome Browser. Metaprofiles were constructed by summing signal intensity over FlyBase r5.49 mRNA annotations.

**Figure 3.**
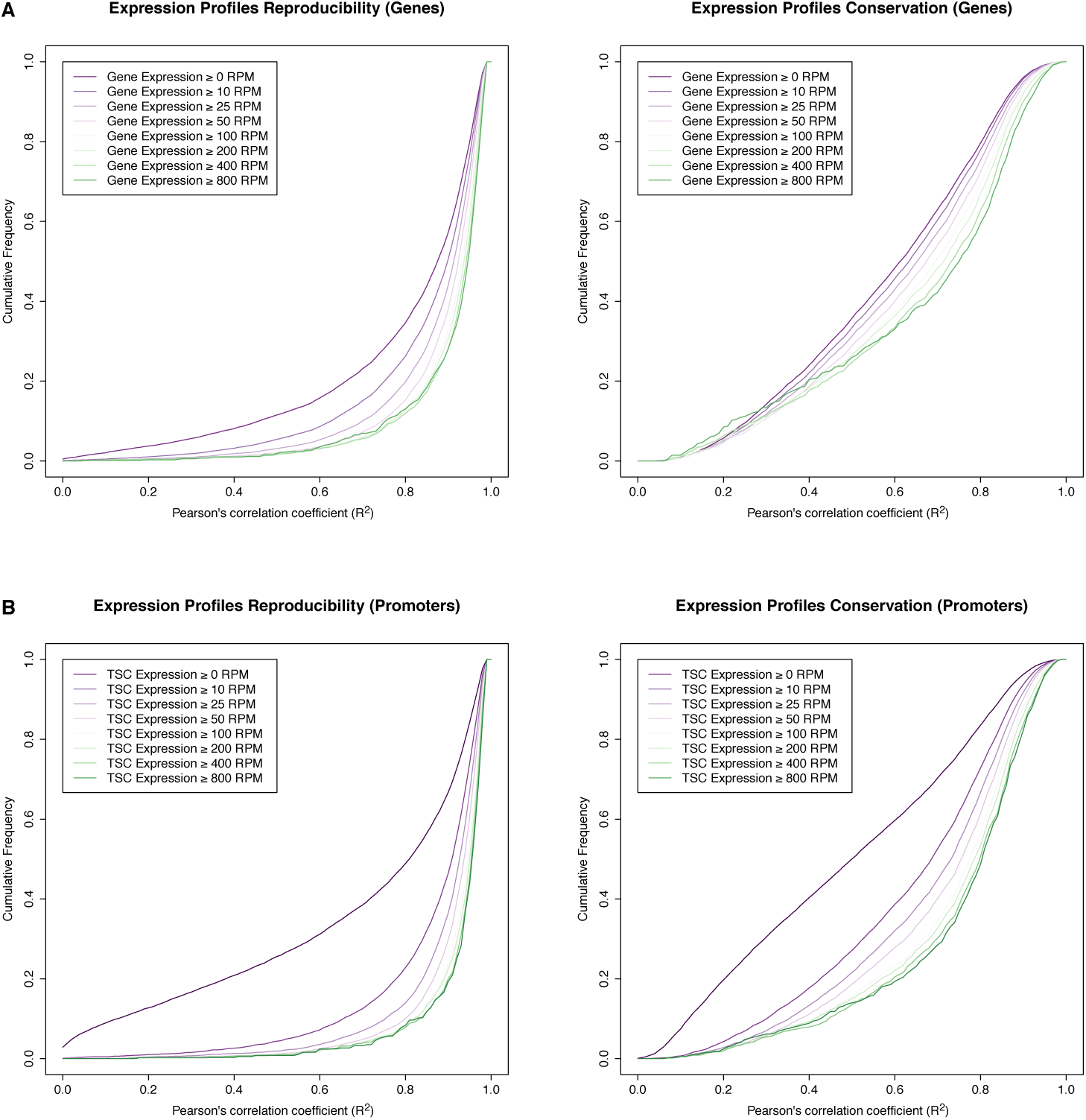
Reproducibility of expression time series. Reproducibility of expression profile measurements for genes (A) and individual TSCs (B). The graphs represent cumulative distributions of expression profile correlations across *D. melanogaster* biological replicates (left), and across all species (right). We only considered genes whose maximum expression level throughout the *D. melanogaster* time series (replicate 1) exceeded a given threshold (see legends; RPM: reads per million). Note that the variation across species vastly exceeds the variation across replicates, at all expression thresholds considered.

**Figure 4.**
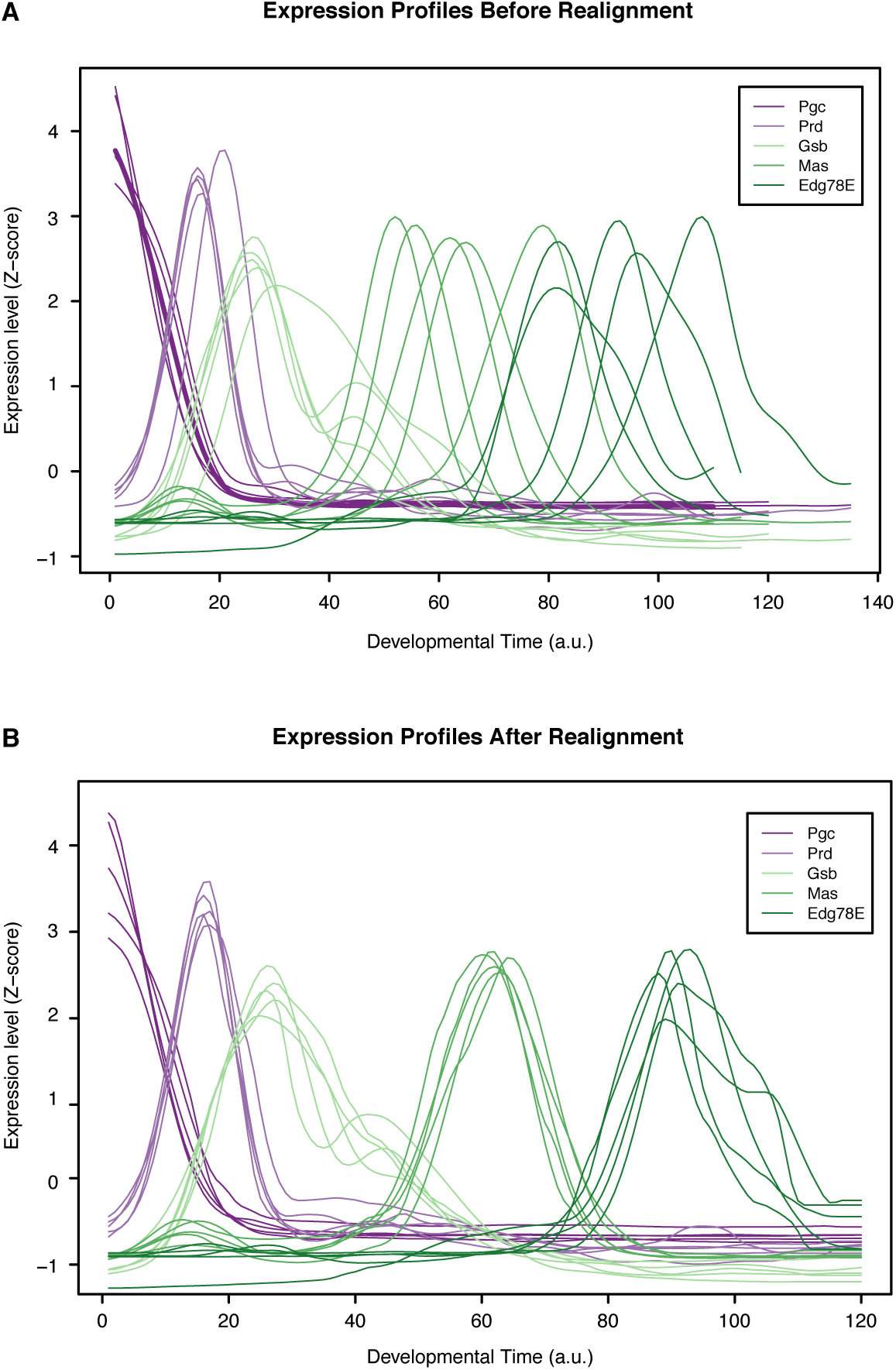
Time series alignment by time-warping of gene expression profiles. Global gene expression profiles from all species were aligned to the *D. melanogaster* time series as described in Methods. This figure shows the expression profiles for well-characterized developmental regulators before (A) and after (B) alignment. The time scale corresponds to the absolute *D. melanogaster* developmental time (24 hours divided into 120 units by upsampling).**B Expression of the *hunchback* Promoters**

**Figure 5.**
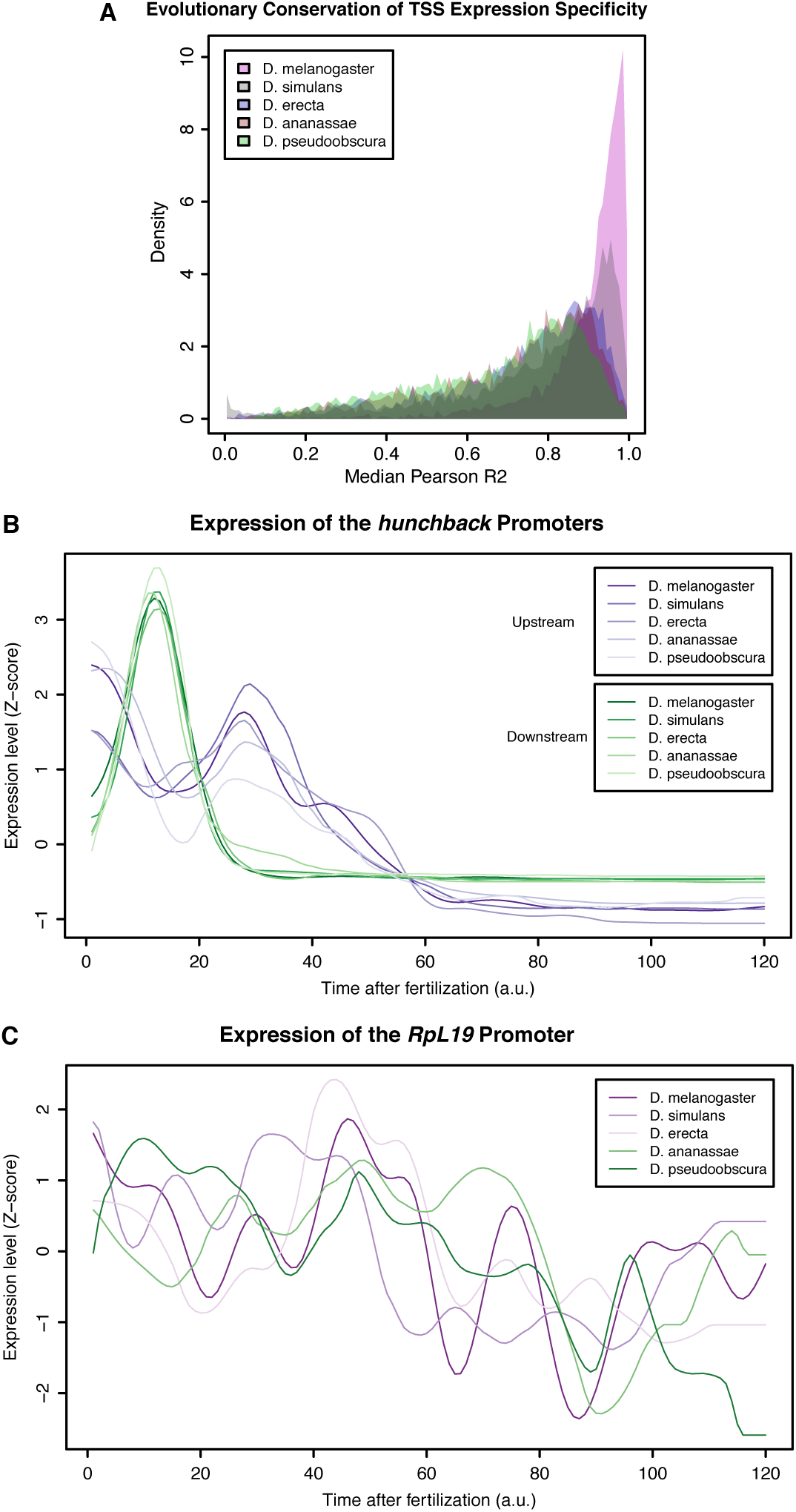
Evolutionary conservation of TSC expression profiles. (A)Distribution of average correlation coefficients for all orthologous TSCs between pairs of species. (B)Aligned expression profiles for the 2 promoters of the *hunchback* gene. (C) Aligned expression profiles for the *RpL19* gene.

**Figure 6.**
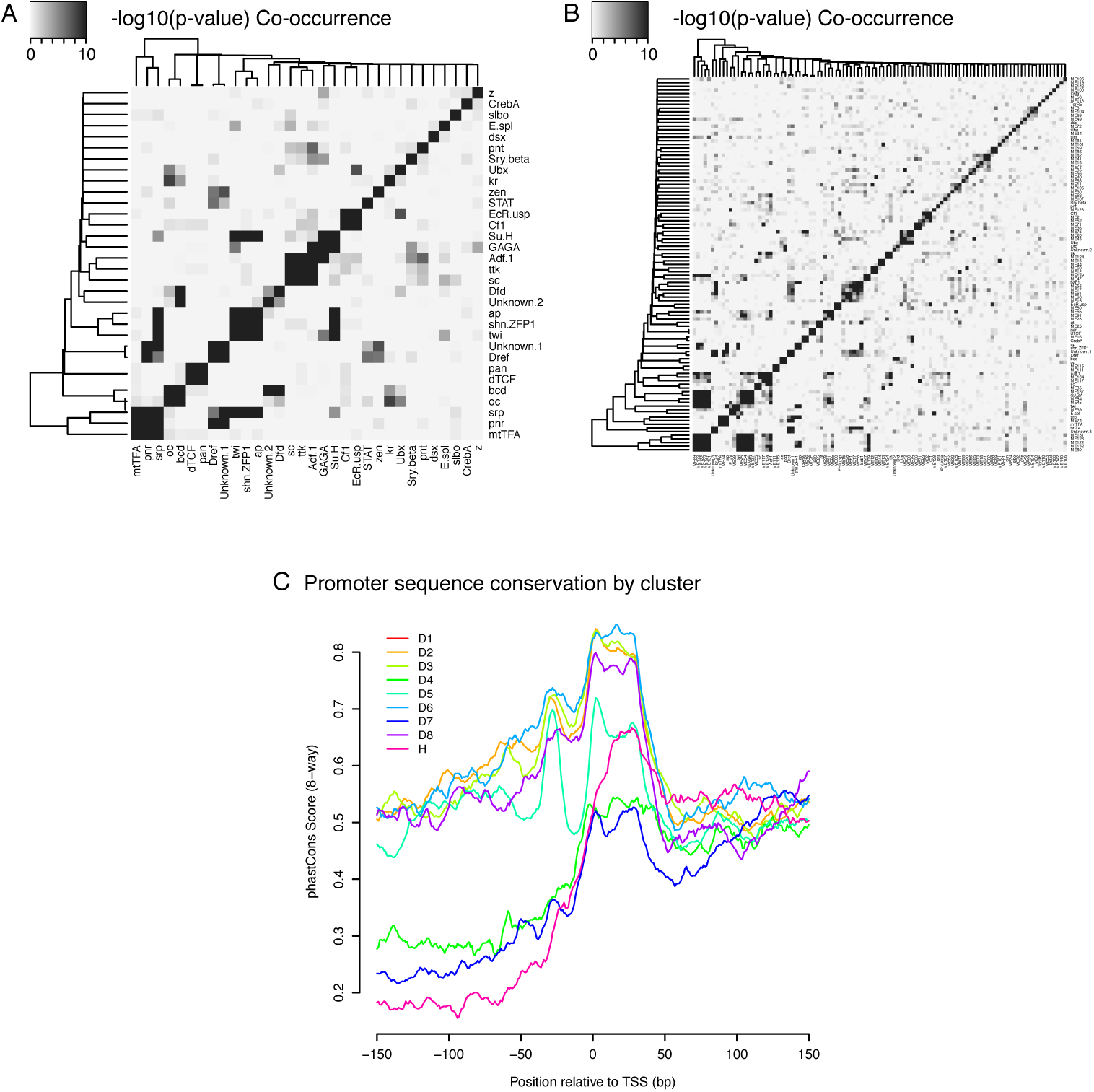
Core promoter types: Co-occurence of TFBSs & Sequence conservation profiles. (A) Co-occurence of TFBSs within the same promoter regions (top 3 motifs for each of the 9 expression clusters) (B) Co-occurence of TFBSs within the same promoter regions (top 10 otifs for each cluster) (C) Median sequence conservation profiles of the promoter regions of the 9 clusters (8-way phastCons scores)

**Figure 7.**
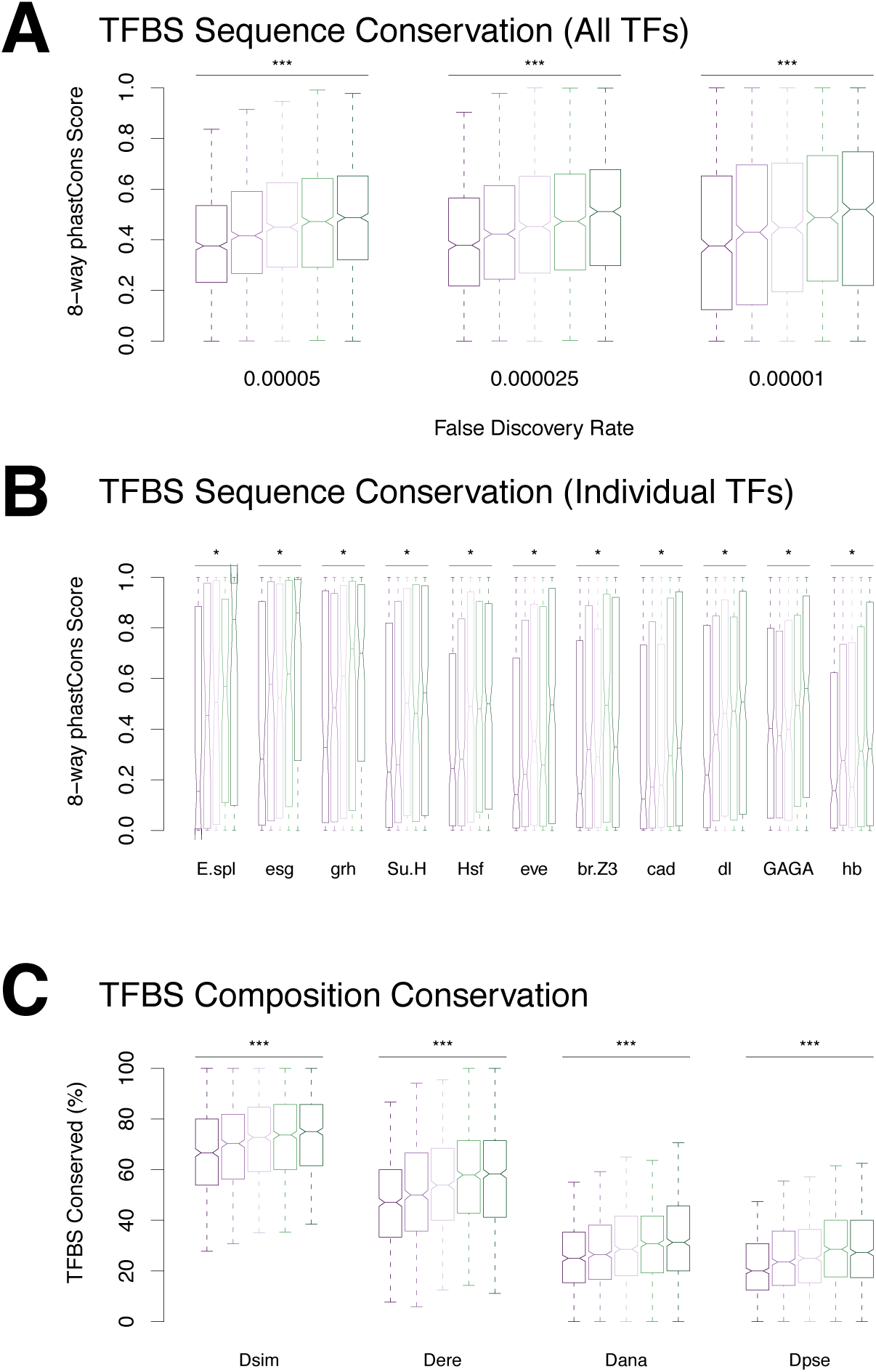
Conservation of ranscription factor binding sites. (A) TFBS sequence conservation versus profile conservation. (B) Conservation of individual motif types. We found a significant correlation between motif sequence conservation and promoter profile conservation for 20 motif types (Bonferroni-corrected p-value < 0.01). (C) Conservation of promoter TFBS composition, regardless of TFBS position.

**Figure 8.**
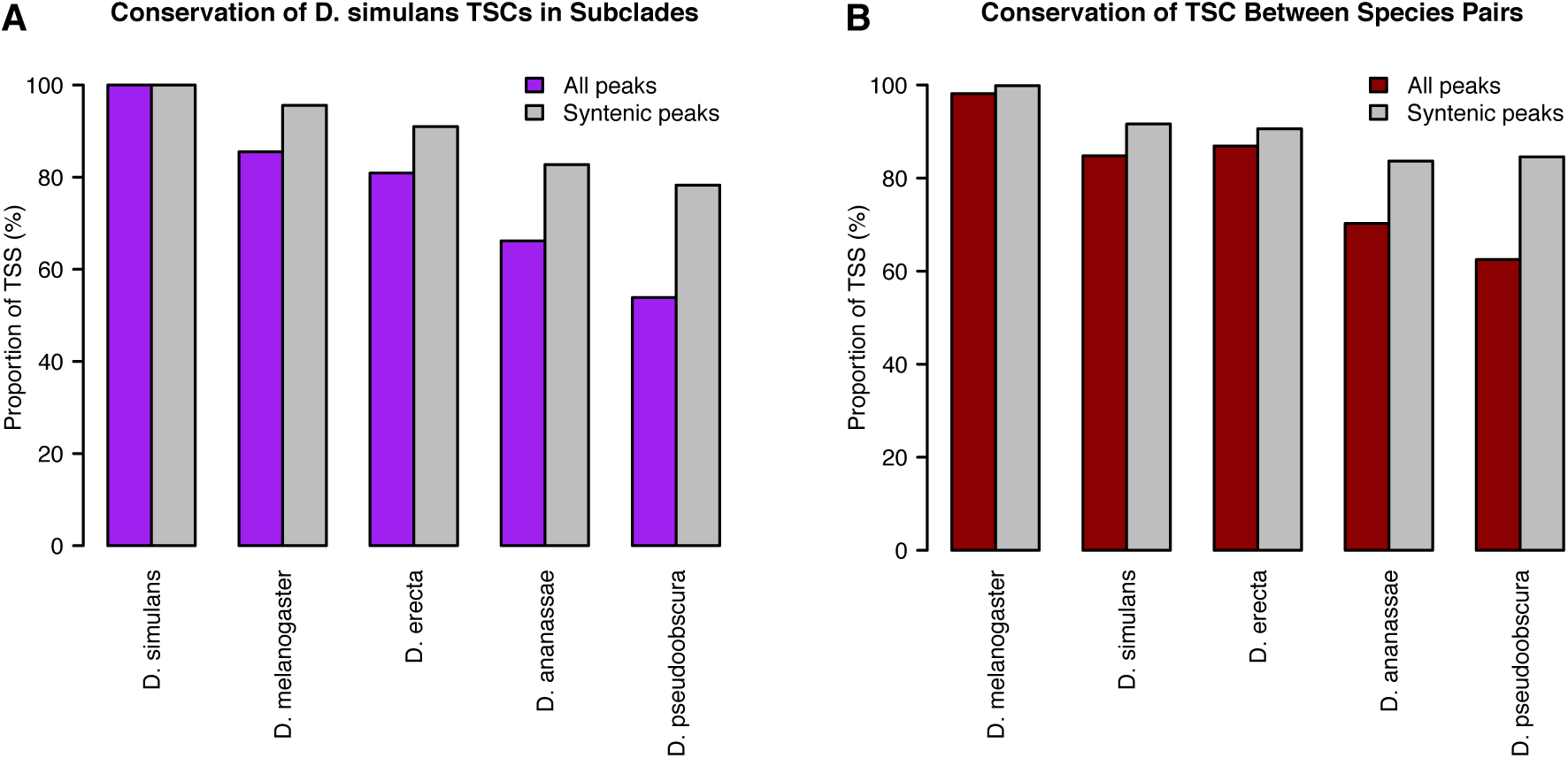
Alternative analyses of TSC conservation. TSC conservation was quantified as described in Methods. (A) *D. simulans-centric* analysis. (B) Quantification of TSC conservation between species pairs, as opposed to subclades.

**Figure 9.**
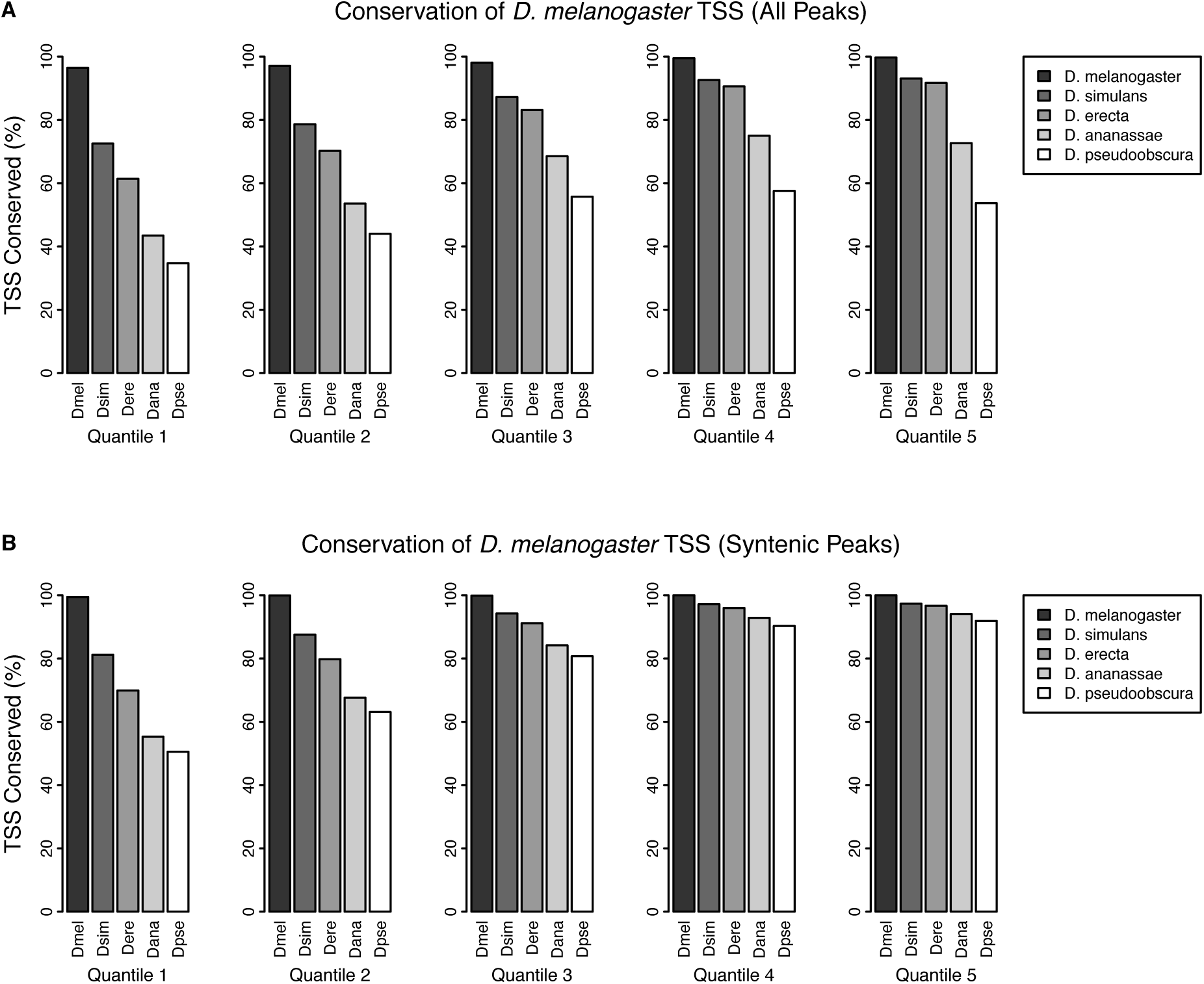
TSC conservation by expression quantiles. *D. melanogaster* TSCs were categorized into 5 expression quantiles based on total raw signal for the full time series. Functional conservation was assessed as described in Methods. (A) Conservation of all TSCs. (B) Conservation of TSCs with syntenic alignments in all 5 species.

**Figure 10.**
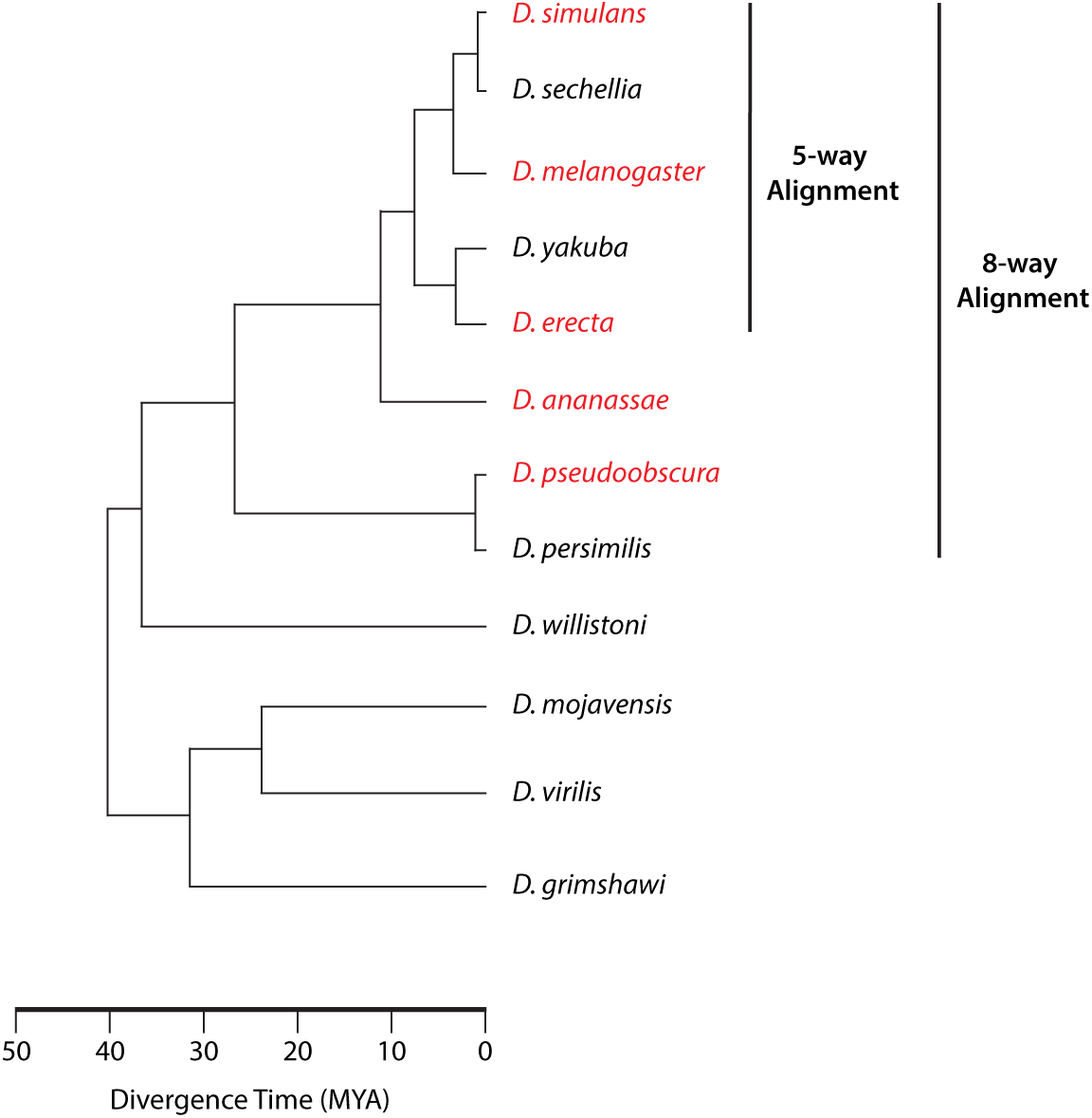
Phylogeny of sequenced species. The species for which we gathered data appear in red. When assessing sequence conservation for features conserved across all 5 species studied, we included the genome sequences of the 5 species studied and the 3 additional sequenced species from the same monophyletic group (8-way alignment). When assessing conservation throughout the *melanogaster* subgroup, we used the genomes of our 3 species and the other 2 from the subgroup (5-way alignment).

**Figure 11.**
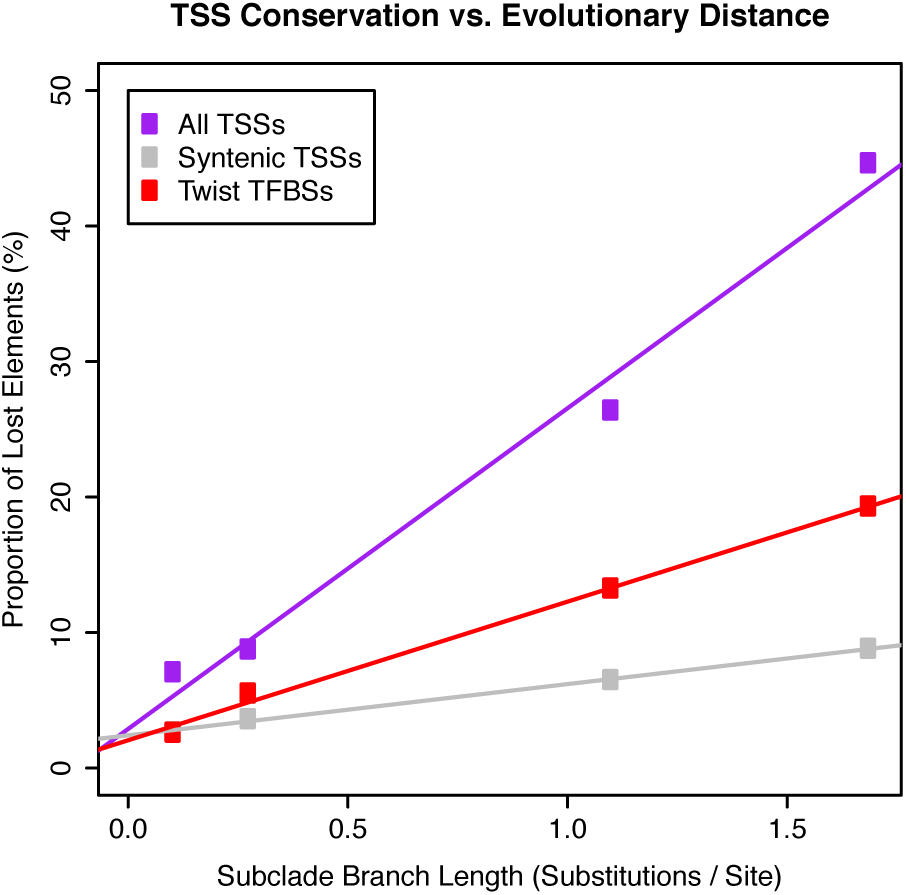
Evolutionary rates of gain and loss for TSCs and Twist TFBSs Twist TFBS data from <Twist paper>.

**Figure 12.**
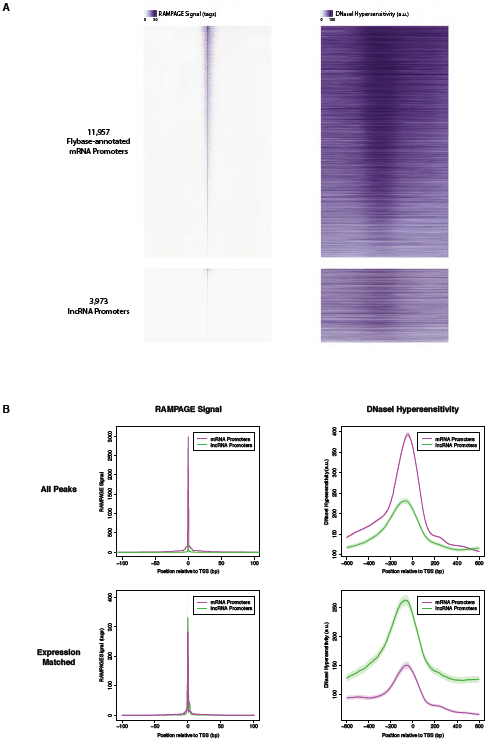
DNase I hypersensitivity at RAMPAGE TSCs. (A) Heatmaps of RAMPAGE signal (left) and DNase-seq signal (right) over individual TSCs. We are comparing TSCs that overlap FlyBase-annotated mRNA transcription start sites (top), which we use as positive controls, to the TSCs of putative lncRNAs (bottom). In each group, TSCs are sorted by total RAMPAGE signal intensity. (B) Class-wise average profiles of RAMPAGE signal (left) and DNase-seq signal (right). The lines represent median profiles, the shaded areas cover +/- 1 standard deviation as estimated by bootstrapping. When considering all peaks (top), lncRNA promoters show weaker DNase sensitivity than FlyBase-annotated controls, but the latter also have considerably stronger RAMPAGE signal. When matching RAMPAGE signal distributions (bottom), this trend is reversed. Shaded areas represent +/-1 standard deviation, as estimated by downsampling and bootstrapping.

**Figure 13.**
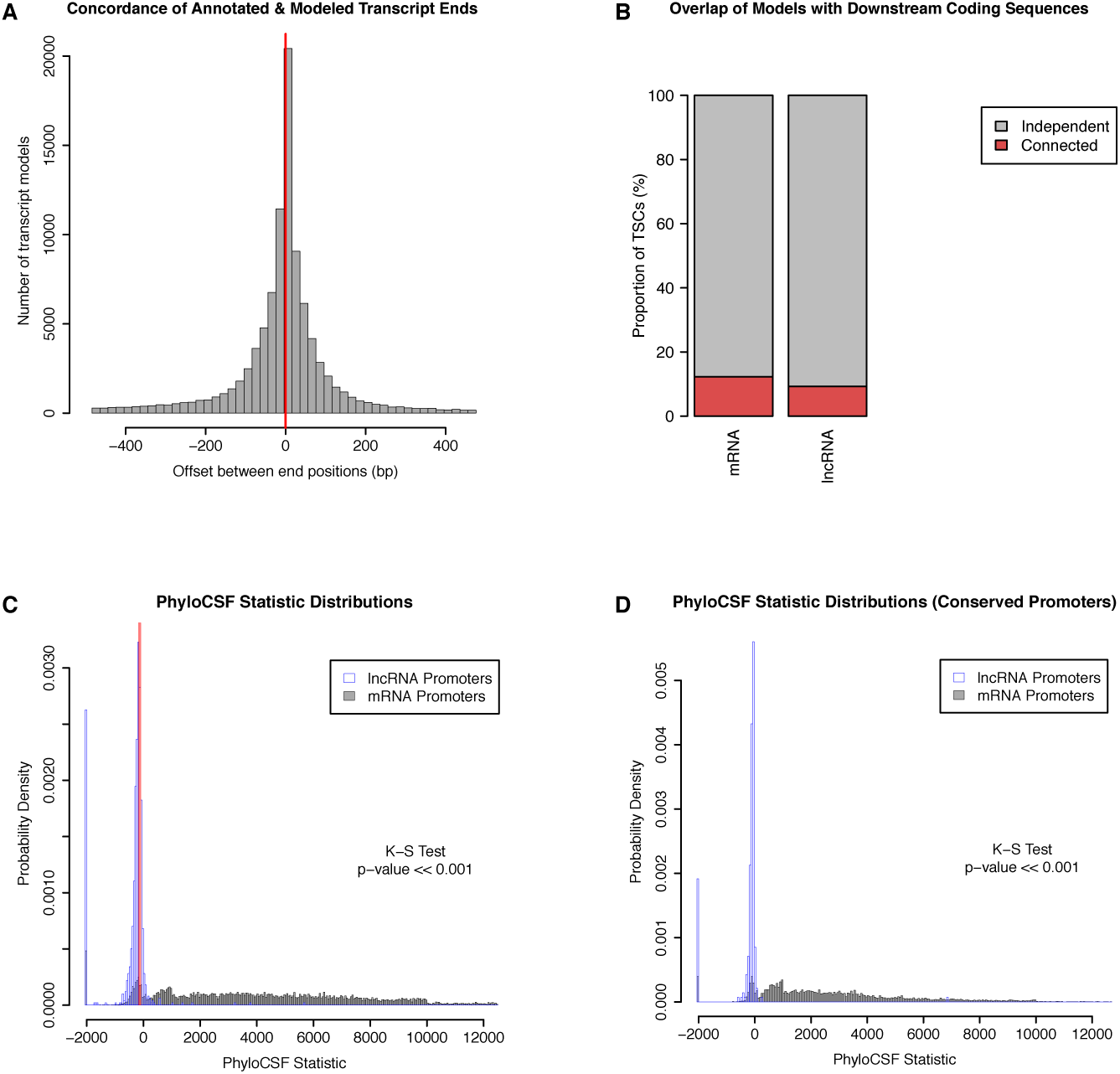
Independence and protein-coding potential of putative lncRNAs. We used published *D. melanogaster* embryo RNA-seq data and Cufflinks to generate transcript models, and only considered those starting within 150bp of a TSC. (A) Concordance of annotated and Cufflinks-modeled 3énds. For each Cufflinks model starting at a protein-coding gene TSC, we are representing the distance to the closest annotated transcript 3énd. (B) Fraction of TSCs for which at least one transcript model overlaps a downstream annotated CDS. Note that while a minority of lncRNA TSC transcript models overlap a downstream protein-coding genes, we observed a similar propensity of Cufflinks models to fuse together consecutive protein-coding genes. (C) PhyloCSF score distributions of protein-coding and putative lncRNA transcript models, as a measure of protein-coding potential. For each TSC, we only considered the transcript model with the highest score. Transcripts with no ORF were assigned a default score of −2,000. Red lines: scores for known lncRNAs (roX1, roX2, bithoraxoid). (D) PhyloCSF analysis restricted to TSCs that are functionally conserved across all 5 species.

**Figure 14.**
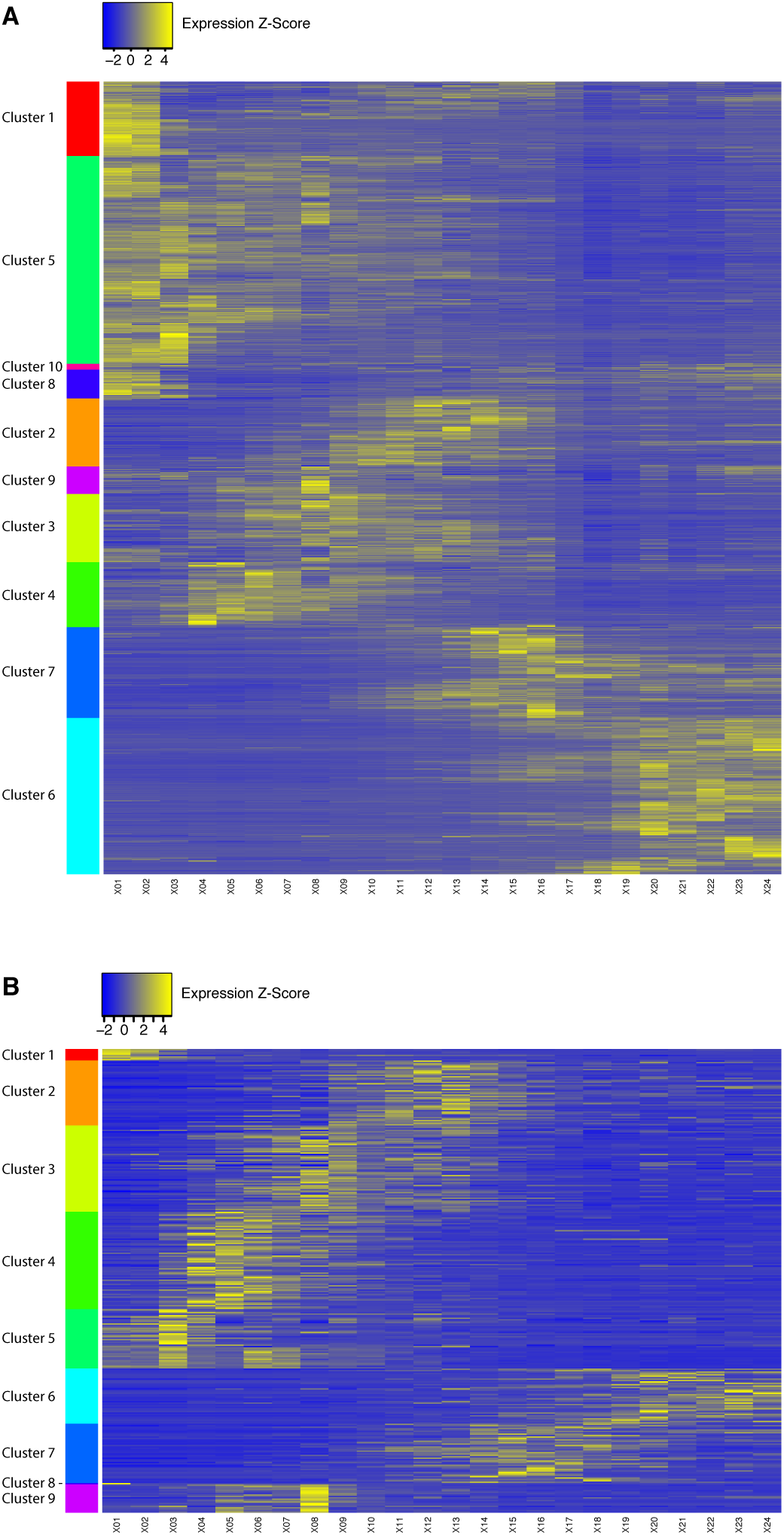
Clustering of *D. melanogaster* developmental expression profiles. In order to analyze coexpression between putative lncRNA and protein-coding gene promoters, we first grouped protein-coding gene promoter expression profiles by k-means clustering (10 clusters) to define reference coexpression sets (A). We then clustered both lncRNA and protein-coging gene promoter profiles together, and extracted the lncRNA promoter profiles corresponding to each expression cluster previously defined (see B).

**Figure 15.**
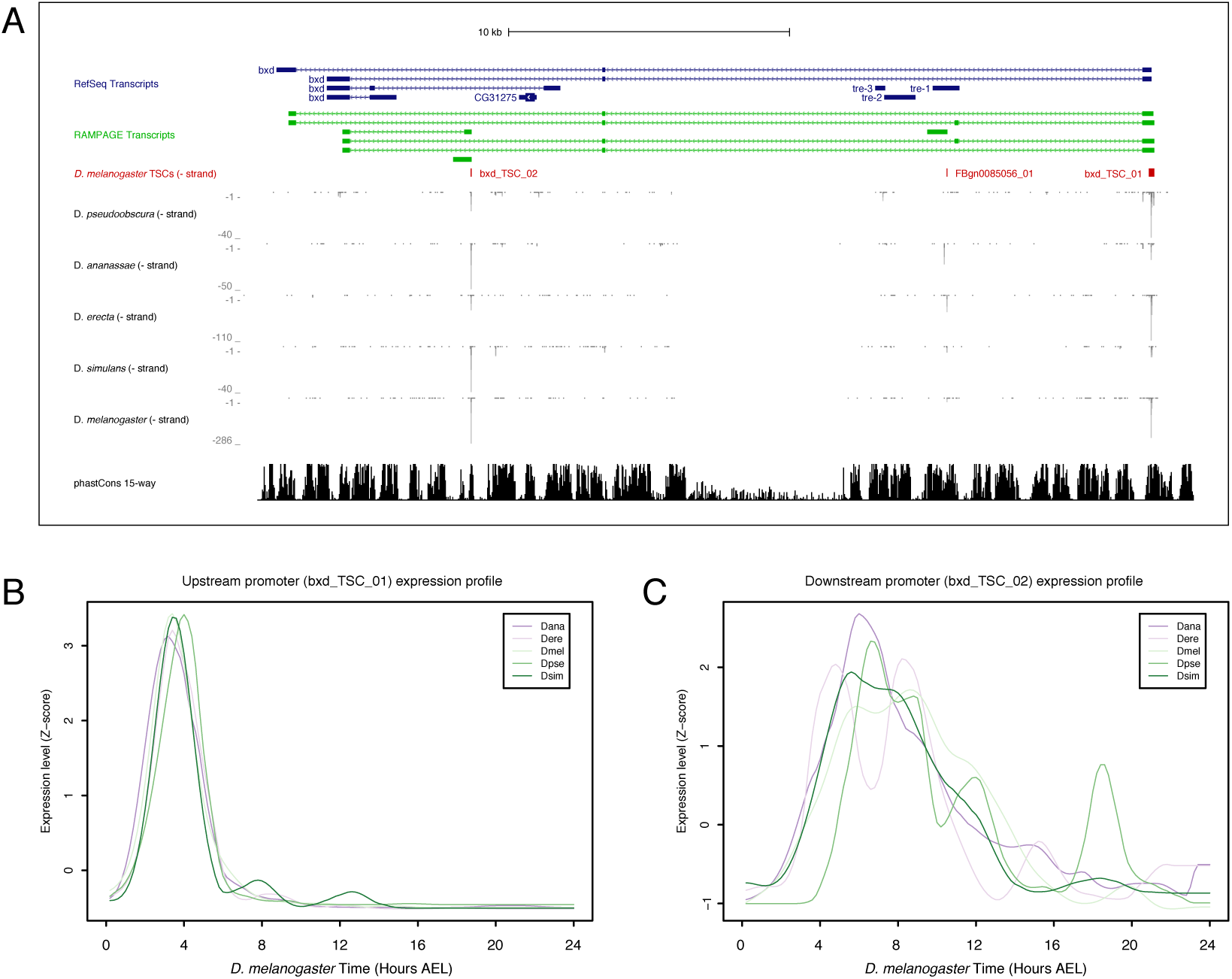
*Bithoraxoid* locus. (A) Organization of the *bithoraxoid* locus (Figure generated with the UCSC Genome Browser). (B) Expression profiles of the upstream *bxd* promoter in the 5 species studied. (C) Expression of the downstream promoter.

**Figure 16.**
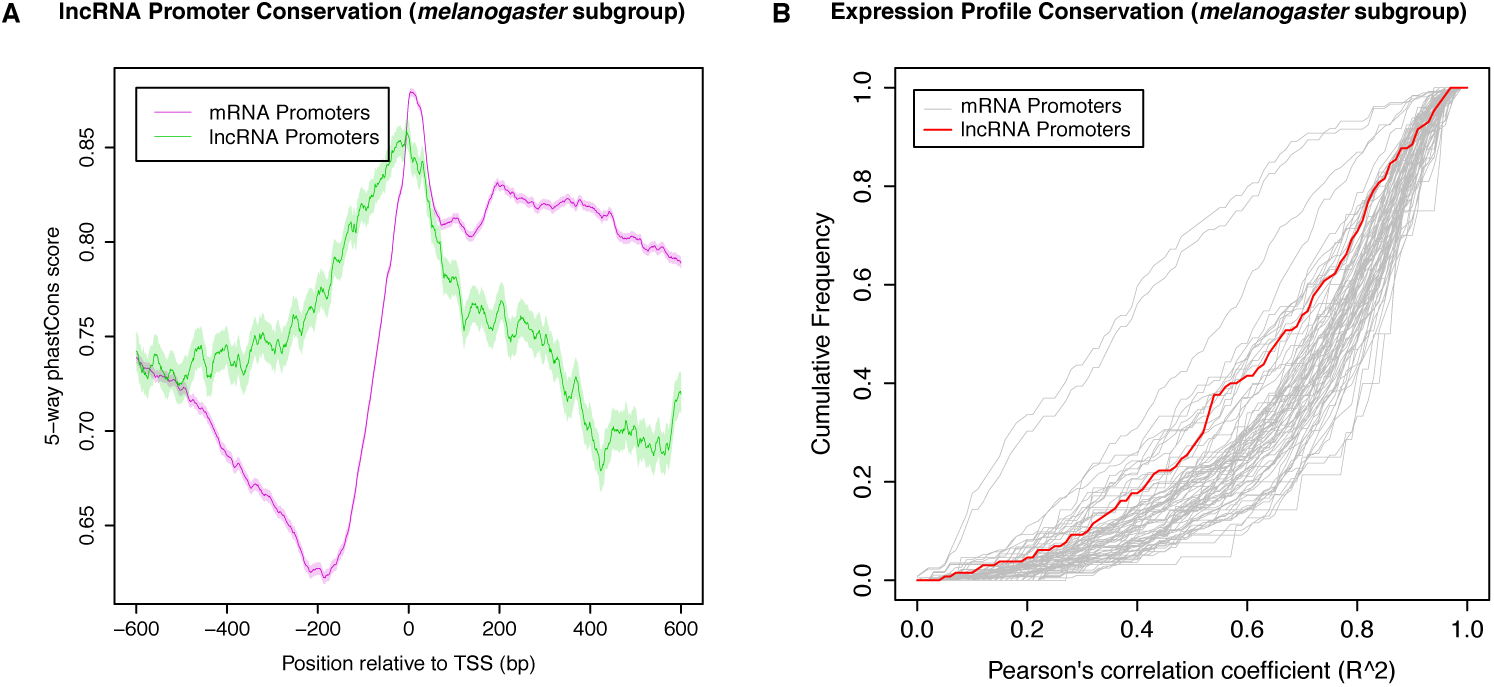
Conservation of *melanogaster* subgroup lncRNA TSCs. (A) Promoter sequence conservation. *melanogaster* subgroup-specific phastCons scores over lncRNA or protein-coding gene promoters that are shared across the 3 species of the subgroup. These phastCons scores were computed by including exclusively the genomes of the 5 sequenced *melanogaster* subgroup species in the input multiple sequence alignment. (B) Expression specificity conservation. All functionally shared promoters with maximum ≥25 RPM and ≥5-fold expression changes in *D. melanogaster* were included in the analysis (130 lncRNA promoters).

**Figure 17.**
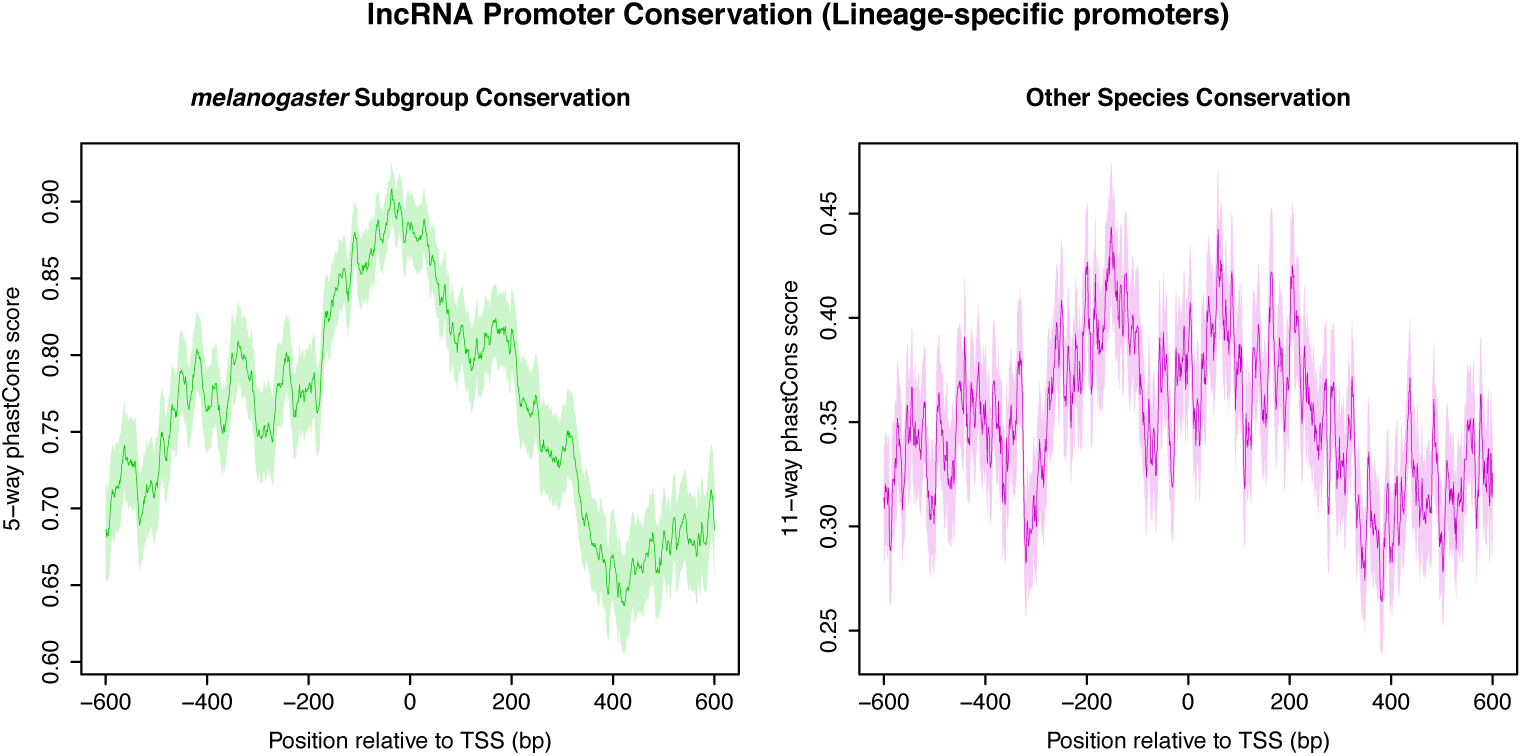
Sequence conservation over *melanogaster* subgroup-specific lncRNA TSCs. (A) Sequence conservation within the *melanogaster* subgroup (5-way phastCons scores). (B) Sequence conservation (phastCons scores based on the alignment of the *D. melanogaster* genome to all outgroup genomes).

**Figure 18.**
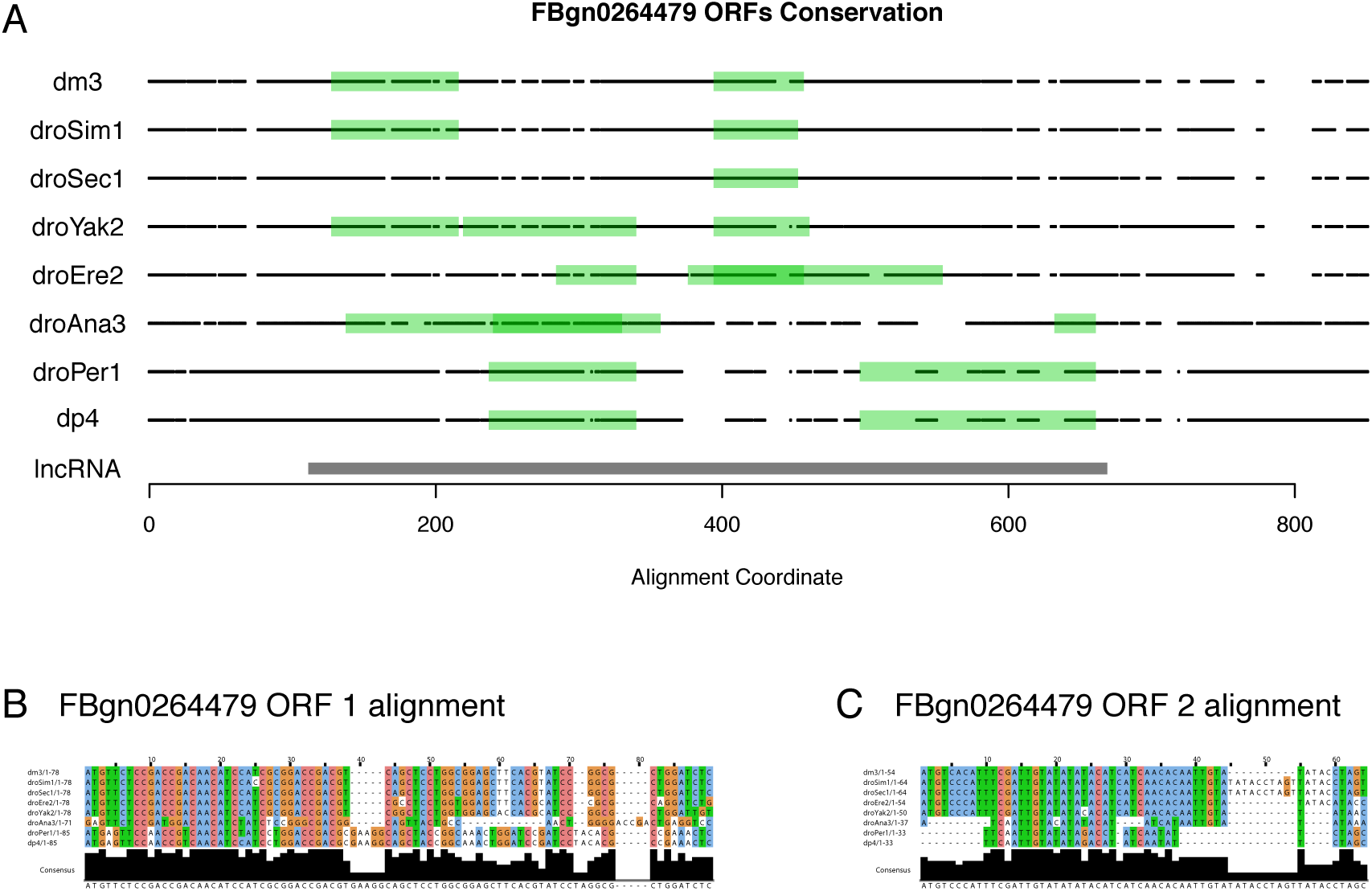
*FBgn0264479* protein-coding potential. (A) Projection of the putative ORFs in all orthologs onto the multiZ multiple sequence alignment (UCSC). Black points: bases aligned to *D. melanogaster* (match or mismatch). White spaces: alignment gaps. lncRNA: *D. melanogaster* transcript. Note the absence of any ORF conserved in all species that express the FBgn0264479 transcript. (B) Multiple sequence alignment for *D. melanogaster* ORF #1. (C) Multiple sequence alignment for *D. melanogaster* ORF #2.

**Figure 19.**
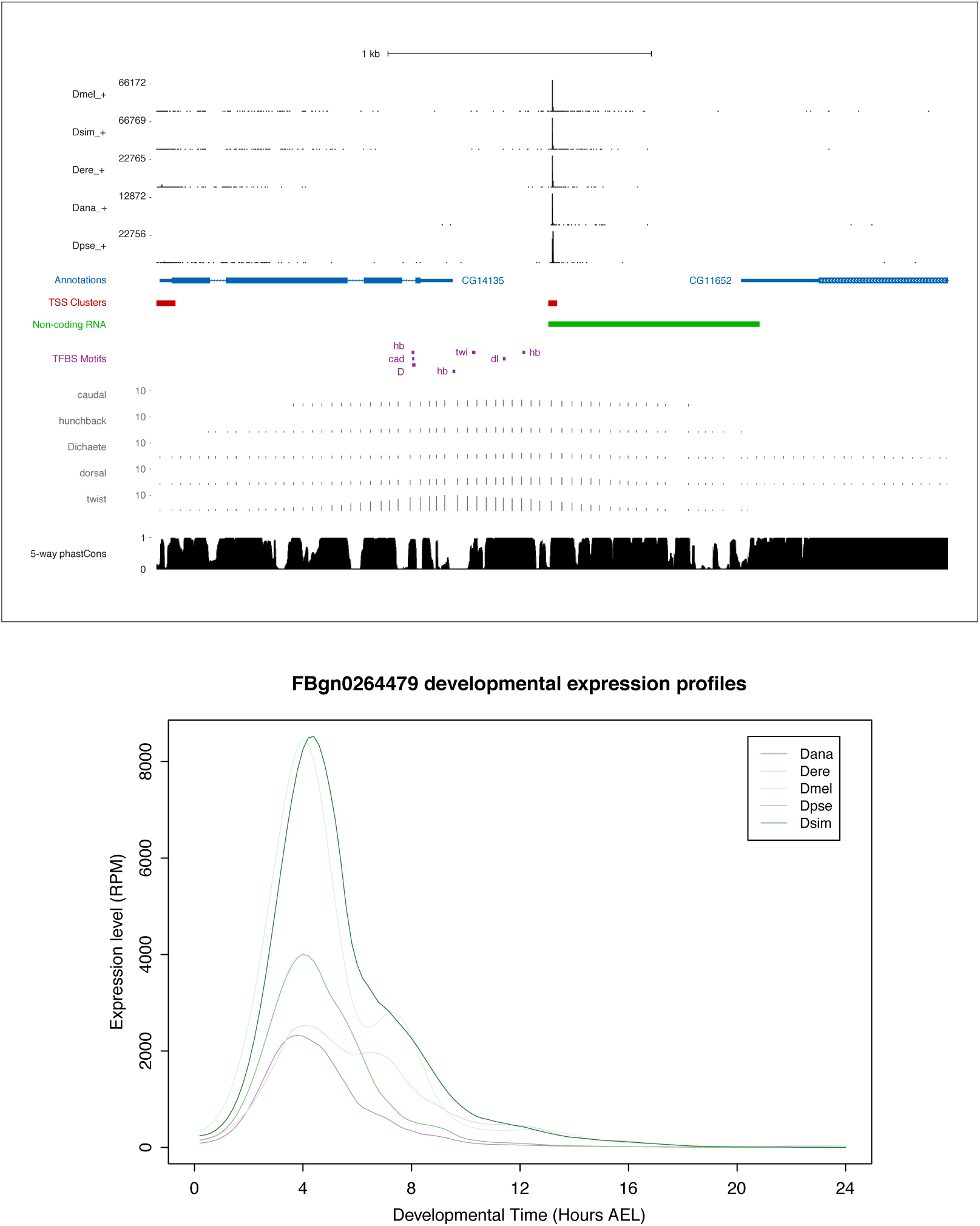
*FBgn0264479* locus and expression. Top: *FBgn0264479* locus organization. Includes all data tracks from Fig. 6A. In addition, the 5 tracks just above the phastCons scores represent the chromatin immunoprecipitation aA§ microarray data (ChIP-chip) for 5 transcription factors: caudal, hunchback, Dichaete, dorsal, twist (modENCODE data visualized on the UCSC Genome Browser). Bottom: Expression profiles in reads per million (RPM).

**Figure 20.**
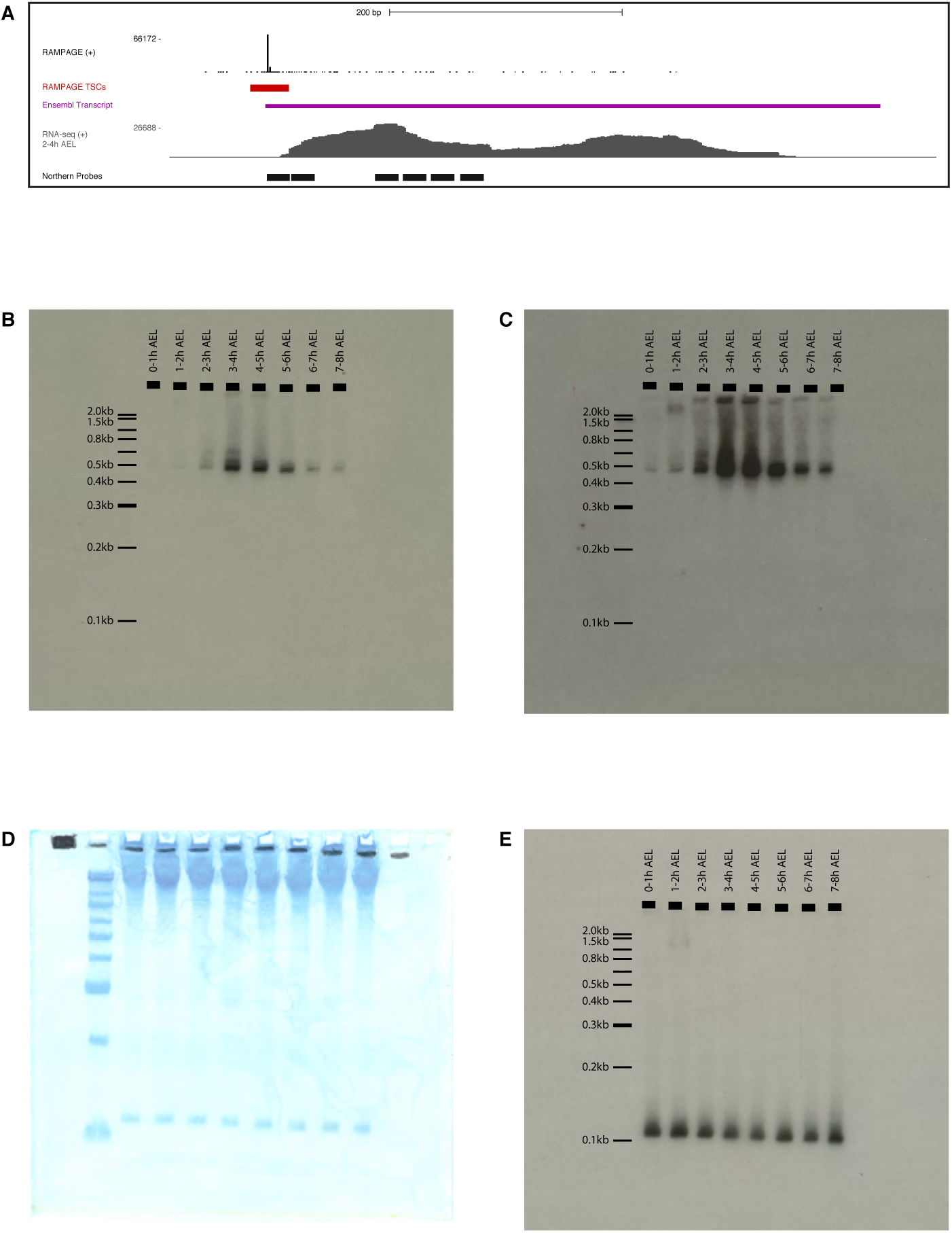
FBgn0264479 Northern-blot. (A) Description of the probes. Tracks from top to bottom: RAMPAGE signal, TSC, Ensembl transcript annotation, RNA-seq signal (modENCODE, 2-4 hours time point), FBgn0264479 probes (antisense to target). We used a mix a all 6 radiolabeled oligonucleotide probes. (B) Exposure of the whole membrane, hybridized with anti-FBgn0264479 probes. (C) Same blot, longer exposure. (D) Methylene blue staining of the membrane prior to hybridization. First lane: Invitrogen 0.1-2.0kb RNA ladder. (E) Hybridization with anti-5S rRNA probe.

**Figure 21.**
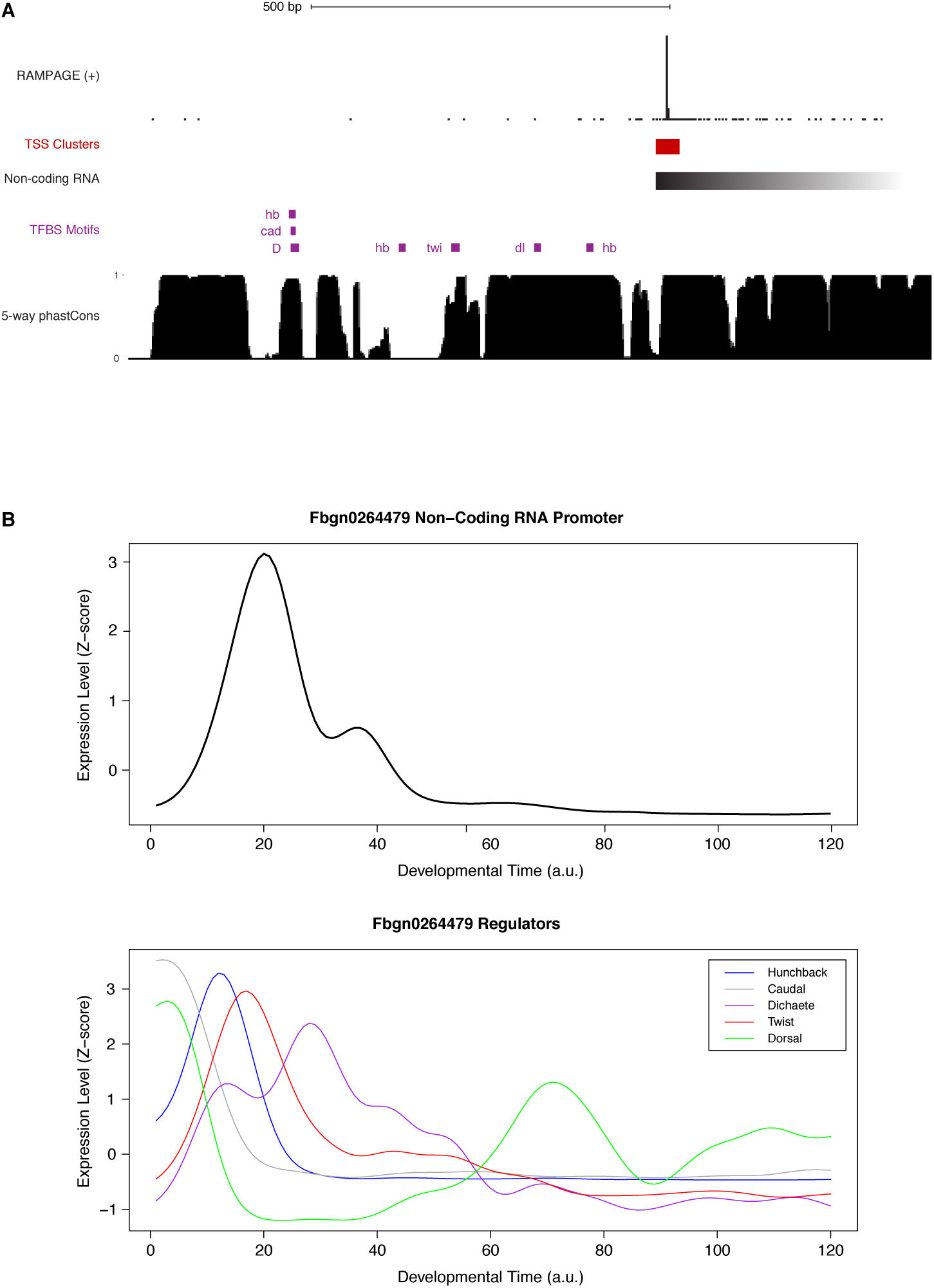
*FBgn0264479* transcriptional regulation. (A) Sequence conservation over the putative binding sites for the factors with ChIP-chip signal over the promoter (see Suppl. Fig. 18). We used FIMO to search for Jaspar-defined motifs within 600bp upstream of the main TSS. Note that all putative TFBSs but one are under strong purifying selection within the *melanogaster* subgroup. (B) Expression profiles of the genes encoding the putative regulators of *FBgn0264479*.

## References

1. Lewis, E.B. A gene complex controlling segmentation in Drosophila. Nature 276, 565–70 (1978).

2. Nusslein-Volhard, C. & Wieschaus, E. Mutations affecting segment number and polarity in Drosophila. Nature 287, 795–801 (1980).

3. Levine, M. & Davidson, E.H. Gene regulatory networks for development. Proc Natl Acad Sci U S A 102, 4936–42 (2005).

4. Peter, I.S. & Davidson, E.H. Evolution of gene regulatory networks controlling body plan development. Cell 144, 970–85 (2011).

5. St Johnston, D. & Nusslein-Volhard, C. The origin of pattern and polarity in the Drosophila embryo. Cell 68, 201–19 (1992).

6. Levine, M. & Hoey, T. Homeobox proteins as sequence-specific transcription factors. Cell 55, 537–40 (1988).

7. Hoey, T. & Levine, M. Divergent homeo box proteins recognize similar DNA sequences in Drosophila. Nature 332, 858–61 (1988).

8. Desplan, C., Theis, J. & O'Farrell, P.H. The sequence specificity of homeodomain-DNA interaction. Cell 54, 1081–90 (1988).

9. Segal, E., Raveh-Sadka, T., Schroeder, M., Unnerstall, U. & Gaul, U. Predicting expression patterns from regulatory sequence in Drosophila segmentation. Nature 451, 535–40 (2008).

10. Banerji, J., Rusconi, S. & Schaffner, W. Expression of a beta-globin gene is enhanced by remote SV40 DNA sequences. Cell 27, 299–308 (1981).

11. Banerji, J., Olson, L. & Schaffner, W. A lymphocyte-specific cellular enhancer is located downstream of the joining region in immunoglobulin heavy chain genes. Cell 33, 729–40 (1983).

12. Zinn, K., DiMaio, D. & Maniatis, T. Identification of two distinct regulatory regions adjacent to the human beta-interferon gene. Cell 34, 865–79 (1983).

13. Small, S., Kraut, R., Hoey, T., Warrior, R. & Levine, M. Transcriptional regulation of a pair-rule stripe in Drosophila. Genes Dev 5, 827–39 (1991).

14. Spitz, F. & Furlong, E.E. Transcription factors: from enhancer binding to developmental control. Nat Rev Genet 13, 613–26 (2012).

15. Schaffner, W. Enhancers, enhancers - from their discovery to today's universe of transcription enhancers. Biol Chem 396, 311–27 (2015).

16. Benoist, C. & Chambon, P. Deletions covering the putative promoter region of early mRNAs of simian virus 40 do not abolish T-antigen expression. Proc Natl Acad Sci U S A 77, 3865–9 (1980).

17. Lenhard, B., Sandelin, A. & Carninci, P. Metazoan promoters: emerging characteristics and insights into transcriptional regulation. Nat Rev Genet 13, 233–45 (2012).

18. Zabidi, M.A. et al. Enhancer-core-promoter specificity separates developmental and housekeeping gene regulation. Nature 518, 556–9 (2015).

19. Lettice, L.A. et al. A long-range Shh enhancer regulates expression in the developing limb and fin and is associated with preaxial polydactyly. Hum Mol Genet 12, 1725–35 (2003).

20. Goodrich, J.A. & Tjian, R. Unexpected roles for core promoter recognition factors in cell-type-specific transcription and gene regulation. Nature Reviews Genetics 11, 549–558 (2010).

21. Gerstein, M.B. et al. What is a gene, post-ENCODE? History and updated definition. Genome Res 17, 669–81 (2007).

22. Gingeras, T.R. Origin of phenotypes: genes and transcripts. Genome Res 17, 682–90 (2007).

23. Kapranov, P. et al. RNA maps reveal new RNA classes and a possible function for pervasive transcription. Science 316, 1484–1488 (2007).

24. Djebali, S. et al. Landscape of transcription in human cells. Nature 489, 101–8 (2012).

25. Graveley, B.R. et al. The developmental transcriptome of Drosophila melanogaster. Nature (2010).

26. Gerstein, M.B. et al. Integrative analysis of the Caenorhabditis elegans genome by the modENCODE project. Science 330, 1775–87 (2010).

27. Young, R.S. et al. Identification and properties of 1,119 candidate lincRNA loci in the Drosophila melanogaster genome. Genome BiolEvol 4, 427–42 (2012).

28. Derrien, T. et al. The GENCODE v7 catalog of human long noncoding RNAs: analysis of their gene structure, evolution, and expression. Genome Res 22, 1775–89 (2012).

29. Augui, S., Nora, E.P. & Heard, E. Regulation of X-chromosome inactivation by the X-inactivation centre. Nat Rev Genet 12, 429–42 (2011).

30. Ulitsky, I. & Bartel, D.P. lincRNAs: genomics, evolution, and mechanisms. Cell 154, 26–46 (2013).

31. Guttman, M. & Rinn, J.L. Modular regulatory principles of large non-coding RNAs. Nature 482, 339–46 (2012).

32. Ponting, C.P., Oliver, P.L. & Reik, W. Evolution and functions of long noncoding RNAs. Cell 136, 629–41 (2009).

33. Tsankov, A., Yanagisawa, Y., Rhind, N., Regev, A. & Rando, O.J. Evolutionary divergence of intrinsic and trans-regulated nucleosome positioning sequences reveals plastic rules for chromatin organization. Genome Res 21, 1851–62 (2011).

34. Schmidt, D. et al. Five-vertebrate ChIP-seq reveals the evolutionary dynamics of transcription factor binding. Science 328, 1036–40 (2010).

35. Stefflova, K. et al. Cooperativity and rapid evolution of cobound transcription factors in closely related mammals. Cell 154, 530–40 (2013).

36. He, Q. et al. High conservation of transcription factor binding and evidence for combinatorial regulation across six Drosophila species. Nat Genet 43, 414–20 (2011).

37. Batut, P., Dobin, A., Plessy, C., Carninci, P. & Gingeras, T.R. High-fidelity promoter profiling reveals widespread alternative promoter usage and transposon-driven developmental gene expression. Genome Res (2012).

38. Carninci, P. et al. Genome-wide analysis of mammalian promoter architecture and evolution. Nature Genetics 38, 626–635 (2006).

39. Valen, E. et al. Genome-wide detection and analysis of hippocampus core promoters using DeepCAGE. Genome Research 19, 255–265 (2009).

40. Hoskins, R.A. et al. Genome-wide analysis of promoter architecture in Drosophila melanogaster. Genome Res (2011).

41. Batut, P. & Gingeras, T.R. RAMPAGE: Promoter Activity Profiling by Paired-End Sequencing of 5'-Complete cDNAs. Curr Protoc Mol Biol 104, 25B 11 1–25B 11 16 (2013).

42. Goltsev, Y. & Papatsenko, D. Time warping of evolutionary distant temporal gene expression data basedon noise suppression. Bmc Bioinformatics 10, - (2009).

43. Kalinka, A.T. et al. Gene expression divergence recapitulates the developmental hourglass model. Nature 468, 811–U102 (2010).

44. FitzGerald, P.C., Sturgill, D., Shyakhtenko, A., Oliver, B. & Vinson, C. Comparative genomics of Drosophila and human core promoters. Genome Biol 7, R53 (2006).

45. Odom, D.T. et al. Tissue-specific transcriptional regulation has diverged significantly between human and mouse. Nature Genetics 39, 730–732 (2007).

46. Villar, D. et al. Enhancer evolution across 20 mammalian species. Cell 160, 554–66 (2015).

47. Haerty, W. & Ponting, C.P. Mutations within lncRNAs are effectively selected against in fruitfly but not in human. Genome Biol 14, R49 (2013).

48. Thomas, S. et al. Dynamic reprogramming of chromatin accessibility during Drosophila embryo development. Genome Biol 12, R43 (2011).

49. Lipshitz, H.D., Peattie, D.A. & Hogness, D.S. Novel transcripts from the Ultrabithorax domain of the bithorax complex. Genes Dev 1, 307–22 (1987).

50. Petruk, S. et al. Transcription of bxd noncoding RNAs promoted by trithorax represses Ubx in cis by transcriptional interference. Cell 127, 1209–21 (2006).

51. Franke, A. & Baker, B.S. The roxl and rox2 RNAs are essential components of the compensasome, which mediates dosage compensation in Drosophila. Mol Cell 4, 117–22 (1999).

52. Li, X.Y. et al. Transcription factors bind thousands of active and inactive regions in the Drosophila blastoderm. PLoS Biol 6, e27 (2008).

53. MacArthur, S. et al. Developmental roles of 21 Drosophila transcription factors are determined by quantitative differences in binding to an overlapping set of thousands of genomic regions. Genome Biol 10, R80 (2009).

54. Arora, K. et al. The Drosophila schnurri gene acts in the Dpp/TGF beta signaling pathway and encodes a transcription factor homologous to the human MBP family. Cell 81, 781–90 (1995).

55. Kutter, C. et al. Rapid turnover of long noncoding RNAs and the evolution of gene expression. PLoS Genet 8, e1002841 (2012).

56. Dai, H. et al. The zinc finger protein schnurri acts as a Smad partner in mediating the transcriptional response to decapentaplegic. Dev Biol 227, 373–87 (2000).

57. Bokel, C., Schwabedissen, A., Entchev, E., Renaud, O. & Gonzalez-Gaitan, M. Sara endosomes and the maintenance of Dpp signaling levels across mitosis. Science 314, 1135–9 (2006).

58. Coumailleau, F., Furthauer, M., Knoblich, J.A. & Gonzalez-Gaitan, M. Directional Delta and Notch trafficking in Sara endosomes during asymmetric cell division. Nature 458, 1051–5 (2009).

59. Gonzalez-Gaitan, M. Signal dispersal and transduction through the endocytic pathway. Nat Rev Mol Cell Biol 4, 213–24 (2003).

60. Fabrowski, P. et al. Tubular endocytosis drives remodelling of the apical surface during epithelial morphogenesis in Drosophila. Nat Commun 4, 2244 (2013).

61. Dobin, A. et al. STAR: ultrafast universal RNA-seq aligner. Bioinformatics 29, 15–21 (2012).

62. Stark, A. et al. Discovery of functional elements in 12 Drosophila genomes using evolutionary signatures. Nature 450, 219–32 (2007).

63. Legendre, F. et al. Whole mount RNA fluorescent in situ hybridization of Drosophila embryos. J Vis Exp, e50057 (2013).

